# Distinct polyamine metabolic signatures underlie differential sensitivity of hippocampal CA1 and CA2 regions to NMDA-induced excitotoxicity

**DOI:** 10.64898/2026.11.20.768153

**Authors:** Aleksandra Skweres, Aleksandra Owczarek, Małgorzata Beręsewicz-Haller, Aleksandra Duchnowska, Martyna Nalepa, Agata Dalka, Mariusz Radkiewicz, Michalina Wypchło, Francesco Bellomi, Claudia Caturano, Aaron B. Bowman, Giorgio Vivacqua, Barbara Zabłocka, Michał Węgrzynowicz

**Affiliations:** Laboratory of Molecular Basis of Neurodegeneration, Mossakowski Medical Research Institute, Polish Academy of Sciences, Warsaw, Poland; Molecular Biology Unit, Mossakowski Medical Research Institute, Polish Academy of Sciences, Warsaw, Poland; Doctoral School of Translational Medicine, Centre of Postgraduate Medical Education, Warsaw, Poland; Laboratory of Mass Spectrometry, Institute of Biochemistry and Biophysics, Polish Academy of Sciences, Warsaw, Poland; Laboratory of Microscopic and Ultrastructural Anatomy, Integrated Research Center PRAAB, Campus Bio-Medico University of Rome, Rome, Italy; School of Health Sciences, Purdue University, West Lafayette, Indiana, USA

**Keywords:** arginase 2, polyamines, hippocampus, region CA2, excitotoxicity, NMDA

## Abstract

Hippocampal regions differ markedly in their susceptibility to excitotoxic injury, although the mechanisms underlying this differential vulnerability remain incompletely understood. Here, we investigated the regional organization of arginine-polyamine metabolism and its contribution to differential excitotoxic vulnerability of CA1 and CA2 pyramidal neurons. Using immunostaining, biochemical assays, and liquid chromatography–mass spectrometry, we characterized pathway-related enzymes and metabolites in rodent hippocampi. We then examined how pharmacological modulation of polyamine synthesis affects NMDA-induced injury in rat organotypic hippocampal cultures.

Expression of arginase 2 (ARG2) protein, an enzyme that provides ornithine for polyamine synthesis, was restricted to CA2 pyramidal neurons in both mouse and rat hippocampus. Arg2 deletion increased hippocampal arginine levels and reduced all three polyamines: putrescine, spermidine, and spermine, supporting a contribution of ARG2 to hippocampal polyamine homeostasis. Analysis of the distribution of other enzymes involved in arginine metabolism, with a focus on polyamine synthesis, revealed that CA2 also exhibited enrichment of ornithine decarboxylase (ODC) and spermine synthase (SMS), together with low neuronal nitric oxide synthase expression. This enzyme profile suggests a metabolic specialization favouring arginine utilization for polyamine synthesis at the cost of nitric oxide production in CA2 pyramidal neurons. NMDA exposure depleted all three polyamines in organotypic cultures without significantly altering related amino acid levels and caused pronounced injury in CA1 but little damage in CA2. ODC inhibition increased NMDA-induced injury in CA2 but not CA1, suggesting that endogenous putrescine synthesis supports CA2 resistance. Conversely, SMS inhibition selectively reduced CA2 injury, and exogenous spermine attenuated this protection, implicating spermine synthesis in CA2 vulnerability. Ifenprodil, an antagonist of GluN2B-containing NMDA receptors, reduced NMDA-induced injury, but spermine completely abolished this protective effect. These findings suggest that spermine-mediated attenuation of the CA2 neuroprotection provided by SMS inhibition cannot be explained primarily by potentiation of GluN2B-containing NMDA receptors during NMDA-induced injury.

Together, these findings identify a previously unrecognized metabolic phenotype of CA2 pyramidal neurons, characterized by selective ARG2 expression and enrichment of polyamine-synthesizing enzymes. This metabolic specialization may contribute to the distinct functional properties of CA2 pyramidal neurons, including their response to NMDA stimulation and excitotoxicity.

## INTRODUCTION

Pyramidal neurons in individual hippocampal subregions exhibit distinct molecular identities, connectivity, physiological properties and response to pathological insults, with particularly pronounced differences between the neighbouring CA1 and CA2 regions (Dudek et al., 2016). While long-term potentiation can be readily induced in CA1 using conventional stimulation protocols, its induction in CA2, particularly at Schaffer collateral inputs, is strongly limited (Zhao et al., 2007). Another important difference involves the susceptibility to injury: CA1 is particularly susceptible, while CA2 is relatively resistant under several pathological conditions, including epilepsy (Blümcke et al., 2013; Schoene-Bake et al., 2014; Steve et al., 2014; Thom et al., 2005), hypoxia/ischemia (Squire et al., 2020), and traumatic brain injury (Kotapka et al., 1992; Maxwell et al., 2003). These contrasting properties make CA1 and CA2 a useful model for investigating the cellular basis of selective neuronal vulnerability, particularly because their characteristic differences can be reproduced in various experimental models *in vivo* (Häussler et al., 2016; Kirino, 1982; T. K. Lee et al., 2022; Nadler et al., 1978; Sadowski et al., 1999; Schmidt-Kastner & Hossmann, 1988; Sloviter, 1983) and *in vitro* (Nadler et al., 1978; Skweres et al., 2026).

Several mechanisms underlying CA2-specific properties have been proposed, although their relationship to resistance to injury remains poorly understood. The CA2-enriched scaffolding protein Regulator of G-protein Signaling 14 (RGS-14) suppresses synaptic potentiation (S. E. Lee et al., 2010), while perineuronal nets surrounding CA2 pyramidal neurons and their excitatory synapses further restrict synaptic plasticity (Carstens et al., 2016). CA2 neurons also exhibit greater calcium-buffering capacity and more efficient calcium extrusion than CA1 neurons, thereby limiting postsynaptic calcium transients (Simons et al., 2009). Furthermore, differences in the expression of mitochondria-associated genes between CA1 and CA2 suggest differences in mitochondrial functions between these subregions (Farris et al., 2019). How these region-specific cellular and molecular properties contribute to the differential vulnerability of CA1 and CA2 to injury remains poorly characterized, although our recent findings suggest that increased activity of the mitochondrial calcium uniporter may contribute to the resistance of CA2 pyramidal neurons to N-methyl-D-aspartate (NMDA)-induced injury (Skweres et al., 2026).

To further explore the mechanisms underlying this differential vulnerability, we investigated whether hippocampal subregions differ in the organization of the arginine (Arg)-polyamine (PA) metabolic pathway and whether PA synthesis contributes to the differential sensitivity of CA1 and CA2 to excitotoxic injury. PAs, including putrescine (Put), spermidine (Spd), and spermine (Spm), are polycationic metabolites that regulate diverse cellular processes, including nucleic acid structure, protein synthesis, ion channel activity, and responses to oxidative stress (Pegg, 2016). Their functions are particularly important in the central nervous system, where they promote autophagy, facilitating the clearance of neurodegeneration-associated protein aggregates (Hofer et al., 2022), support cognitive performance during aging (Hofer et al., 2021), and modulate neurotransmission, particularly glutamatergic transmission mediated by NMDA receptors (NMDARs) (Williams, 1997). PAs are synthesized from ornithine (Orn), an amino acid (AA) produced from Arg by arginase. The hippocampus has been reported to be highly enriched in arginase 2 (ARG2) (Bichell et al., 2017), a mitochondrial arginase isoenzyme whose cellular functions has yet to be fully elucidated and whose expression exhibits restricted regional distribution in the brain (Bichell et al., 2017; Nalepa et al., 2025). However, the detailed distribution of ARG2 and other enzymes involved in PA synthesis within the hippocampus, as well as their potential contribution to differential regional vulnerability to injury, remain unexplored.

We therefore characterized the cellular and subregional distribution of ARG2 and related enzymes in the rodent hippocampus and examined the effects of ARG2 deletion on hippocampal PA and associated AA levels. Using organotypic hippocampal cultures (OHCs), we subsequently assessed NMDA-associated changes in PA homeostasis and investigated how pharmacological targeting of distinct steps in PA synthesis affects NMDA-induced neuronal injury in CA1 and CA2. We found that PA synthesis-related enzymes are strongly enriched in CA2 pyramidal neurons and that modulating this metabolic pathway affects their responses to NMDA excitotoxicity, but has no effect in CA1 pyramidal neurons. These findings identify region-specific organization of PA metabolism as a novel phenotype distinguishing CA1 and CA2 and suggest that differential PA homeostasis, influenced by local PA synthesis capacity, contributes to the distinct properties of pyramidal neurons in these subregions.

## MATERIALS AND METHODS

### Experimental models

#### Animals

The animals were maintained at the Animal Facility of the Mossakowski Medical Research Institute, Polish Academy of Sciences (MMRI PAS), under controlled environmental conditions, with a 12-h light/dark cycle and unrestricted access to food and water. The study used wild-type C57Bl/6J mice (hereafter referred to as Arg2^+/+^), mouse line with deletion of Arg2 gene (STOCK Arg2*^tm1Weo^*/J; RRID:IMSR_JAX:020286; Jackson Laboratory, USA; hereafter referred to as Arg2^-/-^; (Shi et al., 2001)), and Wistar rats. The animals used in the present study had not undergone any experimental procedures *in vivo*. Under the Act of 15 January 2015 on the Protection of Animals Used for Scientific or Educational Purposes, sacrificing animals solely to collect organs or tissues for scientific research does not constitute an experimental procedure (Article 3), and therefore does not require ethical approval, provided that euthanasia is performed by appropriately authorized personnel using a method listed in Annex IV to Directive 2010/63/EU. Gender-balanced groups of the animals were used in all experiments.

Depending on the subsequent experimental use of the collected tissues, animals were euthanized by transcardial perfusion under anaesthesia (adult rats and mice), cervical dislocation (adult mice) or decapitation (rat pups), all of which are methods listed in Annex IV. All procedures were performed by experienced personnel holding certificates issued by the Polish Laboratory Animal Science Association and institutional authorization to perform euthanasia of mice and rats and to collect tissues for experimental purposes.

#### Organotypic hippocampal slice cultures

##### Establishment and maintenance of OHCs

OHCs were prepared as previously described (Skweres et al., 2026). Hippocampi from 7-day-old Wistar rats were sectioned into 400-µm-thick slices using a McIlwain Tissue Chopper (Cavey Laboratory Engineering Co. Ltd., UK). Morphologically appropriate slices were selected under a light microscope (Primo Vert, Zeiss, Germany) and transferred onto Millicell-CM membrane inserts (Millipore, USA). Cultures were maintained at 36°C in a humidified atmosphere containing 5% CO_2_. The culture medium consisted of Neurobasal-A, heat-inactivated horse serum and Hanks’ balanced salt solution, supplemented with B27, HEPES and GlutaMAX (all from Gibco, USA), glucose and Antibiotic-Antimycotic Solution (both from Sigma-Aldrich, USA), at previously described concentrations (Skweres et al., 2026). From days 3 to 7 in vitro (DIV), the serum concentration was gradually reduced and B27 supplementation increased, yielding a serum-free medium containing 2% B27 for the final culture phase.

##### OHC treatment

On DIV8, OHCs were incubated with 2 μM propidium iodide (PI; Sigma-Aldrich) for 30 min. to identify and exclude slices with pre-existing damage. Excitotoxicity was induced by exposure to 25 μM NMDA (Sigma-Aldrich) for 3 h. Ornithine decarboxylase (ODC) was inhibited with difluoromethylornithine (DFMO; 0.2 or 1 mM; Tocris Bioscience), spermine synthase (SMS) with N^1^-cyclohexylpropane-1,3-diamine (APCHA; Fluorochem), and GluN2B-containing NMDARs with ifenprodil (Sigma-Aldrich). Before the main experiments, four independent trial series were performed to assess the effects of APCHA and ifenprodil alone on OHC survival across a range of concentrations, providing the basis for selecting 1 mM APCHA and 15 μM ifenprodil for subsequent experiments (see Supplementary Materials).

DFMO, APCHA, and ifenprodil were added 1 h before NMDA exposure and remained in the medium throughout the subsequent 3-h exposure. In the APCHA experiments, Spm (Pol-Aura, Poland; 100 μM), was applied for 3 h, either alone, with NMDA, or with NMDA and APCHA after 1 h of APCHA preincubation. This delay was intended to allow inhibition of endogenous SMS before exogenous Spm supplementation. In the ifenprodil experiments, Spm was applied for 4 h, either alone throughout, alone for 1 h followed by 3 h with NMDA, or with ifenprodil for 1 h followed by 3 h with ifenprodil and NMDA. This preincubation period allowed exposure to Spm, alone or together with ifenprodil, before the excitotoxic challenge. Each experimental series included appropriate vehicle controls. After treatment, OHCs were maintained for an additional 20 h in fresh culture medium without the experimental compounds, an then incubated with 6 μM PI for 1 h to label damaged neurons and assess neuronal injury.

### Tissue collection and processing

#### Biochemical analyses

Tissue preparation followed published procedures (Nalepa et al., 2025, 2026), with modifications depending on the further analysis. For biochemical analyses, animals were sacrificed by cervical dislocation, and brains, kidneys, and liver were rapidly collected, frozen on dry ice, and stored at -80°C. A single hemisphere per animal was manually cut into 10–12 sections using a scalpel on wet ice, and hippocampi were precisely dissected from each section under a stereomicroscope (SMZ800 N, Nikon, Japan), avoiding contamination with surrounding tissue. For the arginase activity assay, hippocampal tissue was dissected into CA1 and CA2-3 regions, supposed to represent sensitive and relatively resistant areas, respectively. However, once ARG2 expression was found to be restricted to CA2, we discontinued regional dissection because of the small size of this region and the substantial difficulty of accurately identifying its borders in unlabelled tissue. Tissues isolated from all slices of a single animal were combined as one biological replicate, weighed and homogenized in: RIPA buffer (composition described previously; (Nalepa et al., 2025)), supplemented with a protease inhibitor cocktail (Roche, Switzerland) for western blotting; assay buffer XIV from the Arginase Activity Assay Kit (abcam, UK) for arginase activity measurements; PBS for nitrate/nitrite (collectively referred to as NOx) measurements; or 70% ethanol (Stanlab, Poland) for liquid chromatography-mass spectrometry (LC-MS). Further processing is described in the respective analytical procedures below.

#### Microscopic analyses

Animals were deeply anesthetized with pentobarbital (Biowet, Poland) and, once reflexes were absent, were transcardially perfused with PBS (pH 7.4) followed by 4% paraformaldehyde (PFA; Lach-Ner, Czech Republic) solution in PBS. Brains were post-fixed in 4% PFA for 24 h at 4°C and cryoprotected in 30% sucrose (Lach-Ner) in PBS supplemented with 0.1% sodium azide (VWR, USA). Brains were then embedded in PolyFreeze medium (Sigma-Aldrich) and cut into 30-μm-thick sections at -35°C using a Hyrax M 25 microtome (Zeiss) equipped with an MTR freezing unit (SLEE Medical, Germany). Sections were stored in PBS containing 0.1% sodium azide until further processing.

### Western blotting

Western blot was performed according to the previously described protocol (Nalepa et al., 2025). Proteins were separated by SDS-PAGE and transferred onto nitrocellulose membrane using a Mini Trans-Blot Cell system (Bio-Rad, USA). After blocking, membranes were incubated overnight at 16°C with rabbit monoclonal antibodies against ARG1 (Cell Signaling, USA; cat. no. D4E3M) or ARG2 (abcam, UK; cat. no. ab264066), both diluted at 1:1000, followed by a peroxidase-conjugated anti-rabbit secondary antibody (Vector Laboratories, USA; 1:2000). Signal was detected using SuperSignal™ West Dura chemiluminescent substrate (Thermo Scientific, USA) and a FusionX imaging system (Vilber, France). β-actin served as a loading control and was detected after membrane stripping (2% SDS, 0.5 M Tris pH 6.8 and 0.008% β-mercaptoethanol; 1h at 42°C) using a peroxidase-conjugated rabbit anti-β-actin antibody (abcam; cat. no. ab6276; 1:50 000).

### Arginase activity assay

Arginase activity was measured using the Colorimetric Arginase Activity Assay Kit (abcam) following a previously described protocol (Nalepa et al., 2025). Tissue homogenates were adjusted to 4 μg protein/μl, and 20 μl of sample was mixed with 50 μl of reaction mix in a 96-well plate. After pre-incubation at 37°C for 20 min, absorbance was recorded at 570 nm in a kinetic mode for 10 min. using an Infinite M1000 PRO plate reader (Tecan, Switzerland).

### Immunostaining

Hippocampal sections were stained free-floating in 24-well plates on an MR-1 shaker (Biosan, Latvia). Unless otherwise specified, all steps were performed at 16°C. After washing in PBS, sections were subjected to antigen retrieval in 10 mM citrate buffer (Sigma-Aldrich), pH 6.0 at 80°C for 30 min. Nonspecific binding was blocked with 5% normal horse serum (NHS; Vector Laboratories) in PBST (PBS containing 0.5% Tween-20 (Sigma-Aldrich)). For immunohistochemistry (IHC), endogenous peroxidase activity was quenched with methanol (Pol-Aura) and H₂O₂ (Chempur, Poland) in PBST for 1 h; this step was omitted for immunofluorescence (IF). Sections were incubated overnight at 4°C with primary antibodies against neuronal nitric oxide synthase (nNOS; ABclonal, USA; cat. no. A2649), ARG2, ODC1 (ABclonal; cat. no. A3898), spermidine synthase (SRM; ABclonal; cat. no. A8151), or SMS (ABclonal; cat. no. A9380), all diluted 1:200. For double IF, sections were subsequently incubated overnight at 4°C with a mouse antibody against the CA2 pyramidal neuron marker, RGS-14 (BioLegend, USA; cat. no. 821801; 1:200).

For IHC, sections were incubated with a biotinylated anti-rabbit secondary antibody (1:2000) for 3 h, followed by an avidin–peroxidase complex (ABC-HRP kit; Vector Laboratories) for 1 h. The signal was visualized using a DAB substrate kit (Vector Laboratories). Sections were mounted on HistoBond+ slides (Marienfeld, Germany), dried overnight, counterstained with 0.5% cresyl violet (Acros Organics, Belgium) in 400 mM acetate buffer (pH 3.9), dehydrated through graded ethanol (50–100%; Stanlab, Poland), cleared in xylene (Chempur, Poland), and coverslipped with DPX (Merck, Germany).

For IF, sections were incubated for 3 h with Alexa Fluor 555-conjugated anti-rabbit secondary antibody (Invitrogen, USA; 1:2000). For double IF, Alexa Fluor 488-conjugated anti-mouse secondary antibody (Invitrogen; 1:2000) was added to this incubation. All secondary antibodies were diluted in 5% NHS in PBST. IF sections were counterstained with DAPI (1:10 000) in PBS, mounted on glass slides, and coverslipped with FluorSave Reagent (Millipore, USA).

### Microscopy analysis

#### Analysis of immunostained sections

DAB-stained sections were examined using either a CKX41 (Olympus, Japan) or an Axiovert A1 (Zeiss, Germany) bright-field microscopes. Fluorescently stained sections were analyzed using LSM 780 or ELYRA PS.1 confocal microscopy systems (Zeiss) at the Laboratory of Advanced Microscopy Techniques, MMRI PAS.

#### Analysis of PI-labelled OHCs

PI-labelled OHCs were imaged using an LSM 510 confocal microscope at the Laboratory of Advanced Microscopy Techniques, MMRI PAS. Single optical sections were acquired using a 5x objective, excitation/emission wavelengths of 514/550 nm, 11% laser power and a 7.95 AU pinhole. Image analysis was performed by an investigator blinded to the experimental conditions. To assess subregion-specific neuronal damage, PI fluorescence intensity was measured along a manually drawn, 15-pixel-wide line through the middle of the pyramidal cell layer and plotted against distance. CA2 was identified by its position relative to the dentate gyrus, wider pyramidal cell layer, and larger neuronal somata compared with CA1. Pixel intensities were min.-max. normalized to of 0–255, and mean PI intensities in CA1 and CA2 were calculated to compare the extent of damage across the experimental groups. Images were processed using ZEN 3.5 Lite software.

### Nitrate/nitrite assay

Hippocampal NOx levels were measured as previously described (Nalepa et al., 2026). Briefly, PBS homogenates were centrifuged at 18 620 × g for 20 min. at 4°C (Mikro 220R; Hettich, Germany). An aliquot was reserved for protein determination, and the remainder was ultracentrifuged at 120 000 × g for 30 min. at 4°C using (Optima MAX-XP; Beckman, USA). Supernatants were filtered through 10 kDa centrifugal filters. NOx concentrations were determined using the Nitric Oxide Assay Kit (Gibco, USA) according to the manufacturer’s instructions. Nitrates were reduced to nitrites using nitrate reductase for 2 h at 21°C, followed by incubation with 2,3-diaminonaphthalene for 10 min. at room temperature to form fluorescent 1(H)-naphthotriazole. After NaOH addition, fluorescence was measured in white, 96-well plates at excitation/emission wavelengths of 360/430 nm using an Infinite M1000 PRO plate reader. NOx concentrations were calculated from THE same plate standard curve and normalized to protein content.

### LC-MS analysis

Arg, related AAs, and PAs were quantified by LC-MS as previously described (Nalepa et al., 2026). Briefly, homogenized tissue was centrifuged at 14 000 × g for 15 min at 4°C. Pellets were dissolved in 100 mM NaOH + 0.125% Triton X-100 for protein measurements and supernatants were mixed with sodium bicarbonate and isotope-labelled internal standards, derivatized with propionic anhydride/acetonitrile/pyridine, evaporated under nitrogen, and reconstituted in 10% acetonitrile containing 0.1% formic acid. Metabolites were measured using an Acquity UPLC system coupled to a Xevo TQ-S mass spectrometer, with a BEH C18 column and positive electrospray ionization in multiple-reaction-monitoring mode. Quantification used nine-point calibration curves, and data were analyzed in MassLynx v4.2 (Waters, USA) and normalized to protein content.

### Statistics

Comparisons between two independent groups (arginase activity, NOx and LC-MS experiments) were performed using an unpaired t-test. For experiments using OHCs, each culture was considered a biological replicate, whereas individual slices representing the same experimental condition within a culture were considered technical replicates. Values from technical replicates were averaged to obtain a single value for each biological replicate under individual experimental conditions, and these values were used for statistical analysis. The main effects of treatment and hippocampal region, as well as the treatment × region interaction, were assessed using a two-way mixed ANOVA, with repeated measures on the region factor. Treatment was included as a between-subject factor, while region was included as a within-subject factor. Tukey’s post hoc test was used for pairwise comparisons between experimental conditions. All statistical analyses were performed using GraphPad Prism 10.0.0 (GraphPad Software, USA). Data are presented as means ± SEM. A p-value < 0.05 was considered statistically significant.

## RESULTS

### ARG2 accounts for virtually all hippocampal arginase activity while being expressed exclusively in pyramidal neurons of CA2 subregion

Brain arginase is predominantly represented by ARG2, whose distribution remains poorly characterized, but appears to be restricted to selected cell populations. Using a previously validated antibody (Nalepa et al., 2025), further validated in hippocampal homogenates from Arg2^-/-^ mice, we confirmed the previously reported (Bichell et al., 2017) substantial expression of ARG2 protein in the hippocampus of wild-type (Arg2^+/+^) mice (Fig. 1A). ARG1, the other arginase isoenzyme, was undetectable in hippocampal homogenates from both Arg2^+/+^ and Arg2^-/-^ mice (Fig. 1B), indicating the absence of ARG1 expression under control conditions and no compensatory upregulation following ARG2 loss. Arginase catalytic activity was significantly higher in CA2-3 homogenates than in CA1 homogenates from Arg2^+/+^ mice (Fig. 1C), while in Arg2^-/-^ animals, activity was low and did not differ between CA1 and CA2-3 homogenates (Fig. S1A).

**Fig. 1.**
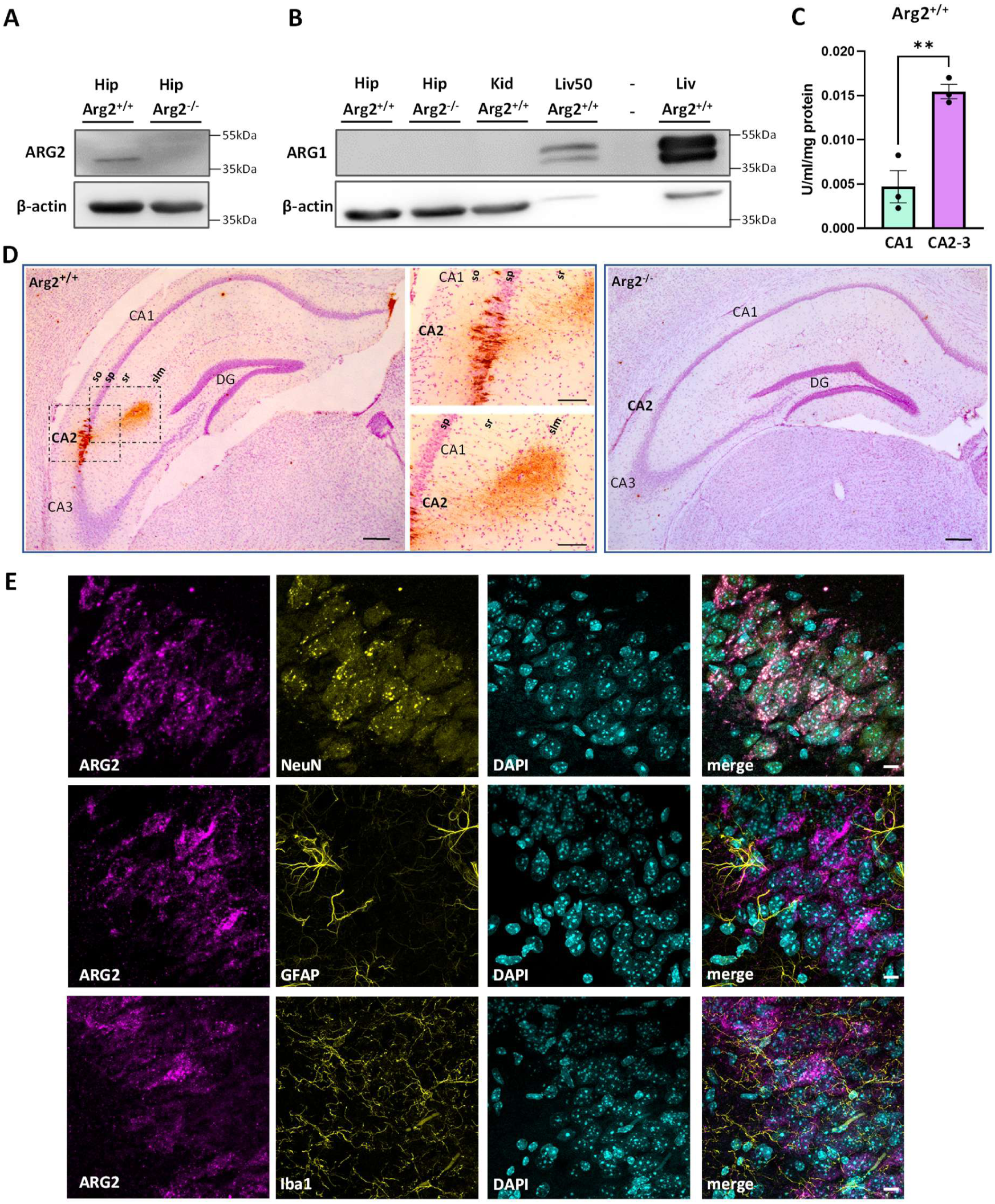
Localization of ARG2 in the mouse hippocampus. **(A)** The anti-ARG2 antibody detects ARG2 protein in the hippocampus of Arg2^+/+^ mice and confirms its complete absence in the hippocampus of Arg2^-/-^ mice. β-actin was used as a loading control. The positions of the molecular weight markers are indicated on the right side of the blot. **(B)** ARG1 was not detected in the hippocampus of either Arg2^+/+^ or Arg2^-/-^ mice. Antibody specificity was assessed using liver tissue, where ARG1 is highly abundant, and kidney tissue, which lacks ARG1, but is enriched in ARG2. β-actin was used as a loading control. The positions of the molecular weight markers are indicated on the right side of the blot. **(C)** Arginase catalytic activity was measured in homogenates of CA2-3 and CA1 hippocampal regions of Arg2^+/+^ mice. Significantly higher activity was observed in CA2-3 than in CA1 (0.0155 ± 0.0008 units/ml/mg of protein vs. 0.0047 ± 0.0018, respectively); p = 0.0057; n = 3 biological replicates (individual animals)). Data are presented as mean ± SEM; **p < 0.01, unpaired t-test. **(D)** ARG2 immunohistochemistry in the hippocampus of Arg2^+/+^ mice reveals strong enrichment in the CA2 region (left panel). ARG2 localizes to the cell bodies of CA2 pyramidal neurons (upper middle panel; enlargement of the corresponding area outlined by a dashed line in the left panel) and their processes, particularly to the slm (lower middle panel; enlargement of the corresponding area outlined by a dashed line in the left panel). Other hippocampal regions show little to no detectable ARG2 staining. No ARG2 signal is detected in the hippocampus of Arg2^-/-^ mice (right panel) confirming the specificity of the staining. All sections were counterstained with Nissl stain (purple). Representative images are shown. Scale bars: 200 µm (left and right panels) and 100 µm (middle panels). **(E)** Co-immunofluorescence staining confirms the neuronal localization of ARG2 in CA2. NeuN, GFAP, and Iba1 were used as markers of neurons, astrocytes, and microglia, respectively. ARG2 is detected in NeuN-positive cells in the pyramidal cell layer but not in GFAP- or Iba1-positive cells. Representative images are shown. Scale bars: 10 µm. Abbreviations: ARG1-arginase 1 protein; ARG2 – arginase 2 protein; Arg2^+/+^ – wild type, Arg2-positive mice; Arg2^-/-^ – Arg2 knockout mice; DG – dentate gyrus; GFAP – glial fibrillary acidic protein, astrocytic marker; Hip – hippocampus; Iba1– ionized calcium-binding adaptor molecule 1, microglial marker; Kid – kidney; Liv – liver; Liv50 – liver tissue homogenate diluted 50x relative to brain samples; NeuN – neuronal nuclear antigen, neuronal marker; slm – stratum lacunosum moleculare; so – stratum oriens; sp – stratum pyramidale; sr – stratum radiatum.

Immunohistochemical analysis further revealed the regional distribution of ARG2. ARG2 labelling was predominantly observed in neuronal somata in the stratum pyramidale (sp) and distal apical dendritic neuropil in the stratum lacunosum-moleculare (slm), with weaker labelling in the proximal apical dendritic neuropil in the stratum radiatum (sr) and little detectable labelling in the basal dendritic compartment of the stratum oriens (so), in both mice (Fig. 1D) and rats (Fig. S1B). No evident somatic labelling was observed in other hippocampal subregions (Fig. 1D, Fig S1B), and no signal was detected in the hippocampus of Arg2^-/-^ mice (Fig. 1D), supporting selective ARG2 expression in hippocampal CA2 pyramidal neurons. The neuron-specific localization of ARG2 in CA2 was confirmed by its presence in NeuN-positive cells, as well as its absence from GFAP-labelled astrocytes and Iba1-labelled microglia (Fig. 1E). Together, these findings identify selective ARG2 expression and associated arginase activity as a novel metabolic phenotype of CA2 pyramidal neurons.

### ARG2 is essential for maintaining physiological PAs levels in the hippocampus

The two major roles attributed to arginase are to compete with nitric oxide synthase (NOS) isoenzymes for arginine (Arg), thereby limiting nitric oxide (NO) production, and to supply Orn for PAs biosynthesis (Fig. 2A). We therefore measured metabolites associated with both pathways in the hippocampus of Arg2^-/-^ mice. Hippocampal levels of nitrite and nitrate (NO breakdown products, collectively denoted as NOx) did not differ between Arg2^+/+^ and Arg2^-/-^ mice (Fig. 2B). Similarly, levels of citrulline (Cit), a second product of NOS activity (Fig. 2A), remained unchanged (Fig. 2C). Meanwhile, Arg levels in hippocampal homogenates from Arg2^-/-^ mice were significantly elevated (Fig. 2C). These findings suggest that greater Arg availability following ARG2 loss does not stimulate NO production in hippocampus.

**Fig. 2.**
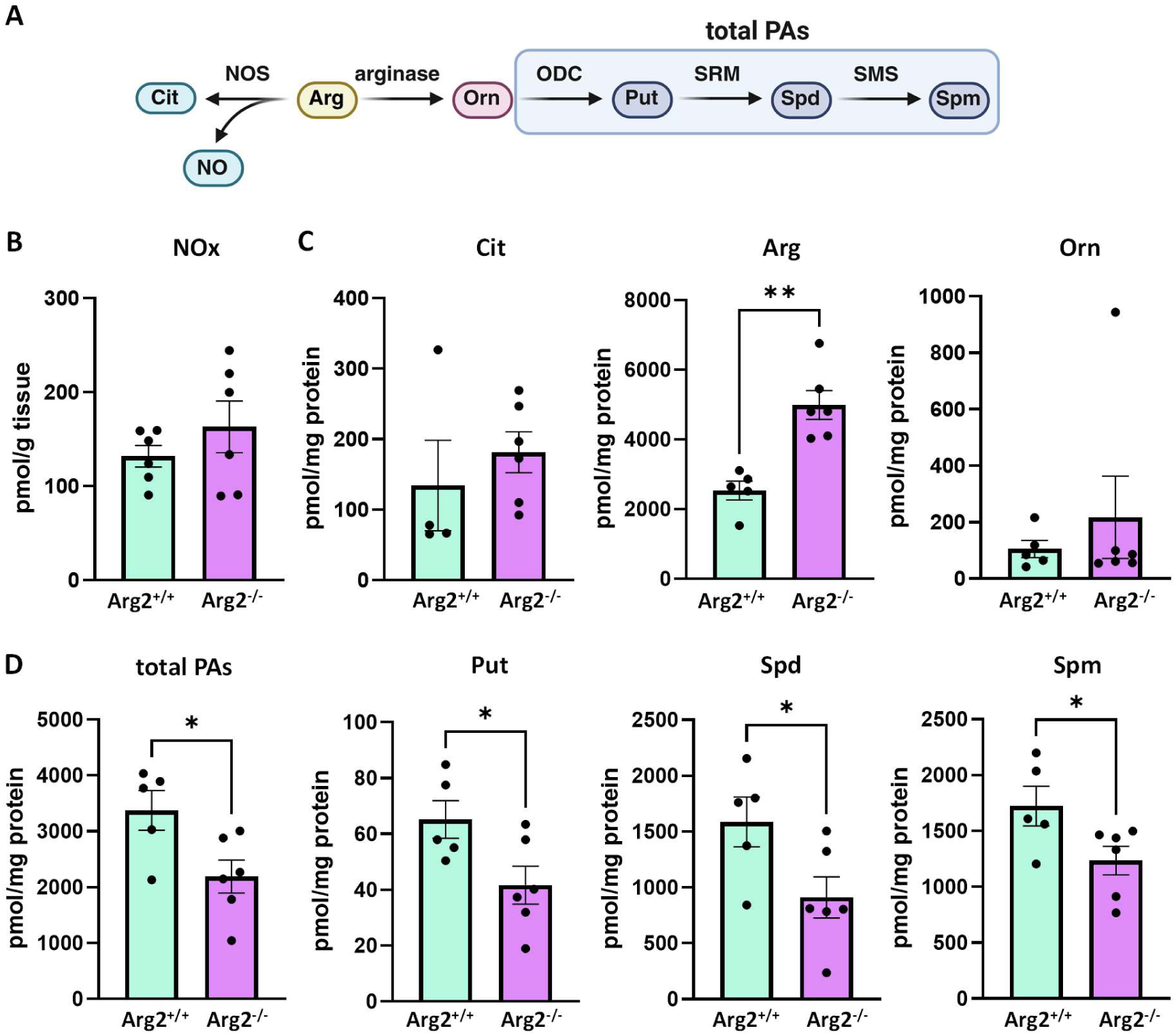
Effect of ARG2 loss on levels of PAs and related AAs in the mouse hippocampus. **(A)** Schematic representation of Arg metabolic pathway. **(B)** Measurements of NOx in the hippocampi of Arg2^+/+^ and Arg2^-/-^ mice. No statistically significant differences in NOx levels were observed between Arg2^+/+^ and Arg2^-/-^ mice (131.83 ± 11.57 pmol/g tissue vs. 163.05 ± 27.47 pmol/g tissue, respectively; p = 0.311; n = 6 biological replicates (individual animals)). **(C)** Measurements of Arg and related AA levels in the hippocampi of Arg2^+/+^ and Arg2^-/-^ mice. No statistically significant differences in Cit levels were observed between Arg2^+/+^ and Arg2^-/-^ mice (134.31 ± 64.21 pmol/mg protein vs. 181.44 ± 29.04 pmol/mg protein, respectively; p = 0.472; n = 4–6 biological replicates (individual animals)). A statistically significant increase in Arg levels was observed in Arg2^-/-^ compared to Arg2^+/+^ mice (4991.10 ± 412.92 pmol/mg protein vs. 2532.31 ± 269.77 pmol/mg protein, respectively; p = 0.001; n = 5–6 biological replicates (individual animals)). No statistically significant differences in Orn levels were observed between Arg2^+/+^ and Arg2^-/-^ mice (105.15 ± 30.44 pmol/mg protein vs. 217.01 ± 145.60 pmol/mg protein, respectively; p = 0.511; n = 5–6 biological replicates (individual animals)). **(D)** Measurements of PA levels in the hippocampi of Arg2^+/+^ and Arg2^-/-^ mice. A statistically significant decrease in total PAs levels was observed in Arg2^-/-^ compared to Arg2^+/+^ mice (2188.44 ± 296.25 pmol/mg protein vs 3374.08 ± 355.55 pmol/mg protein, respectively; p = 0.029; n = 5–6 biological replicates (individual animals)). A statistically significant decrease in Put levels was observed in Arg2^-/-^ compared to Arg2^+/+^ mice (41.67 ± 6.76 pmol/mg protein vs 65.18 ± 6.75 pmol/mg protein, respectively; p = 0.037; n = 5–6 biological replicates (individual animals)). A statistically significant decrease in Spd levels was observed in Arg2^-/-^ compared to Arg2^+/+^ mice (911.07 ± 184.04 pmol/mg protein vs. 1586.69 ± 223.50 pmol/mg protein, respectively; p = 0.043; n = 5–6 biological replicates (individual animals)). A statistically significant decrease in Spm levels was observed in Arg2^-/-^ compared to Arg2^+/+^ mice (1235.71 ± 128.49 pmol/mg protein vs. 1722.21 ± 177.81 pmol/mg protein, respectively; p = 0.049; n = 5-6 biological replicates (individual animals)). Data are presented as mean ± SEM. *p < 0.05, **p < 0.01, unpaired t-test. Abbreviations: AA/AAs – amino acid/amino acids; Arg – arginine; ARG2 – arginase 2; Arg2^+/+^ – wild type, Arg2-positive mice; Arg2^-/-^ – Arg2 knockout mice; Cit – citrulline; NO – nitric oxide; NOS – nitric oxide synthase; NOx – nitrates + nitrites; markers of nitric oxide production; ODC – ornithine decarboxylase; Orn – ornithine; PA/PAs – polyamine/polyamines; Put – putrescine; SMS – spermine synthase; Spd – spermidine; Spm – spermine; SRM – spermidine synthase.

We next examined Orn, the product of the arginase reaction, and its downstream metabolites, PAs (Fig. 2A). Hippocampal Orn levels remained unchanged in Arg2^-/-^ mice (Fig. 2C), possibly reflecting compensation by alternative metabolic pathways. In contrast, total Pas levels were reduced (Fig. 2D), with all three PAs showing significant decreases of similar extent (Fig. 2D), which left their ratios statistically unaltered (Fig. S2A). Levels of the PAs degradation intermediates, N^1^-acetylspermidine (N^1^-AcSpd) and N^1^-acetylspermine (N^1^-AcSpm) did not differ between Arg2^+/+^ and Arg2^-/-^ hippocampus (Fig. S2B), suggesting that ARG2 loss does not affect breakdown of PAs.

These results support a conclusion that ARG2 supports hippocampal PAs synthesis, even though Orn levels remain unchanged following Arg2 deletion. Given that ARG2 expression within the hippocampus is restricted to pyramidal neurons of CA2 (Fig. 1D, E), these results implicate this subregion as an important site of hippocampal PAs production.

### CA2 exhibits enrichment of PAs synthesis enzymes

Having demonstrated a heterogeneous distribution of ARG2 within the hippocampus and a significant reduction in hippocampal PAs levels in Arg2^-/-^ mice, we examined the spatial distribution of enzymes involved across the entire PAs synthesis pathway in mouse hippocampus. To delineate their distribution relative to CA2, we co-immunolabelled each enzyme with RGS-14, an established marker of CA2 pyramidal neurons (S. E. Lee et al., 2010). ARG2 localized exclusively to RGS-14-positive cells and processes and was virtually absent from RGS-14-negative structures (Fig. 3A, Fig. S3A), confirming its restricted localization to CA2 pyramidal neurons. ODC, which converts Orn to Put (Fig. 2A) was strongly enriched in CA2 pyramidal somata, with relatively high expression observed also in proximal CA3 (CA3a and CA3b subregions), but near-background staining in CA1 (Fig. 3B. Fig. S3B). SRM, responsible for Spd synthesis from Put (Fig. 2A) was strongly expressed throughout the whole sp, with little, if any, CA2 enrichment (Fig. 3C. Fig. S3C). SMS involved in conversion of Spd to Spm (Fig. 2A) was clearly enriched in CA2, particularly in pyramidal cell nuclei, with visibly weaker staining in CA1 and CA3 somata (Fig. 3D. Fig. S3D). While ARG2 was detected in CA2 pyramidal somata in sp, and dendrites in sr, and especially slm (Fig. 1D, Fig. 3A, Fig. S3A), the other enzymes were predominantly somatic with no apparent dendritic staining (Fig. 3B-D, Fig. S3B-D). Similar distributions of ODC, SRM and SMS were observed in the rat hippocampus (Fig. S4A-C), suggesting cross-species conservation of these regional patterns.

**Fig. 3.**
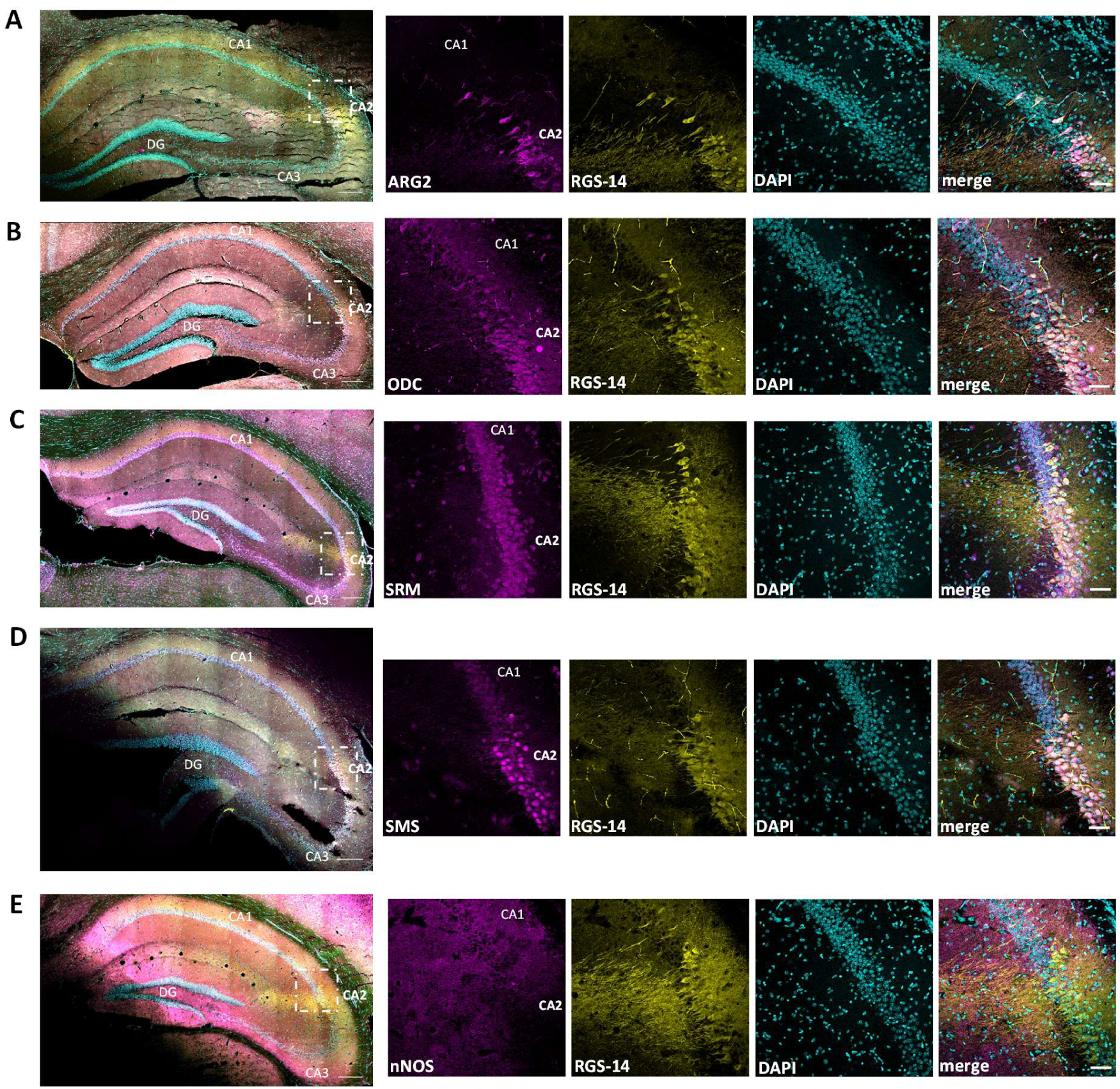
Distribution of PA synthesis-related enzymes in the mouse hippocampus. Localization of **(A)** ARG2, **(B)** ODC, **(C)** SRM, **(D)** SMS and **(E)** nNOS was assessed in mouse hippocampus using immunofluorescence (magenta). For each protein examined, the slices were co-immunostained for RGS-14, a specific marker of CA2 pyramidal neurons (yellow) and counterstained with DAPI (cyan). **(A–E)** Left panels show merged-channel overview images of the entire hippocampus; dashed boxes indicate the CA1/CA2 border region shown at higher magnification in the adjacent single-channel and merged panels. Representative images are shown. Scale bars: 200 µm (overviews), 50 µm (magnified panels). Abbreviations: ARG2 – arginase 2; DAPI – 4′,6-diamidino-2-phenylindole; fluorescent nuclear stain; DG – dentate gyrus; nNOS – neuronal nitric oxide synthase; ODC – ornithine decarboxylase; PA/PAs – polyamine/polyamines; RGS-14 – Regulator of G-protein Signaling 14; marker of CA2 pyramidal neurons; SMS – spermine synthase; SRM – spermidine synthase.

Given the established competition between arginase and NOS for Arg and our finding that NO production markers nevertheless remained unchanged in Arg2^-/-^ mouse hippocampus (Fig. 2B, C), we examined the distribution of neuronal NOS isoenzyme (nNOS) across hippocampal subregions. This analysis revealed noticeably lower nNOS staining intensity in CA2 pyramidal somata than in those of CA1 and CA3 (Fig. 3E, Fig. S3E), potentially explaining why ARG2 loss and the resulting hippocampal Arg accumulation did not alter NO production markers.

Together, these findings reveal pronounced subregional heterogeneity in neuronal Arg-metabolizing enzyme expression in the hippocampus, with a particularly distinctive profile in CA2. CA2 pyramidal neurons show strong enrichment of PAs-synthesizing enzymes alongside low nNOS expression, defining a previously unrecognized metabolic phenotype of these neurons that favours Arg-to-PAs metabolism over NO synthesis relative to neighbouring CA3 and, especially, CA1 neurons.

### NMDA exposure results in reduced PAs levels in OHCs

To explore the functional significance of these findings, we employed a rat OHC-based *in vitro* model of NMDA-induced excitotoxicity (Skweres et al., 2026). First, we measured Arg, related AAs and PAs in slice homogenates from vehicle- and NMDA-treated cultures, to assess the response of the hippocampal PAs system (Fig. 4A) to excitotoxicity.

**Fig. 4.**
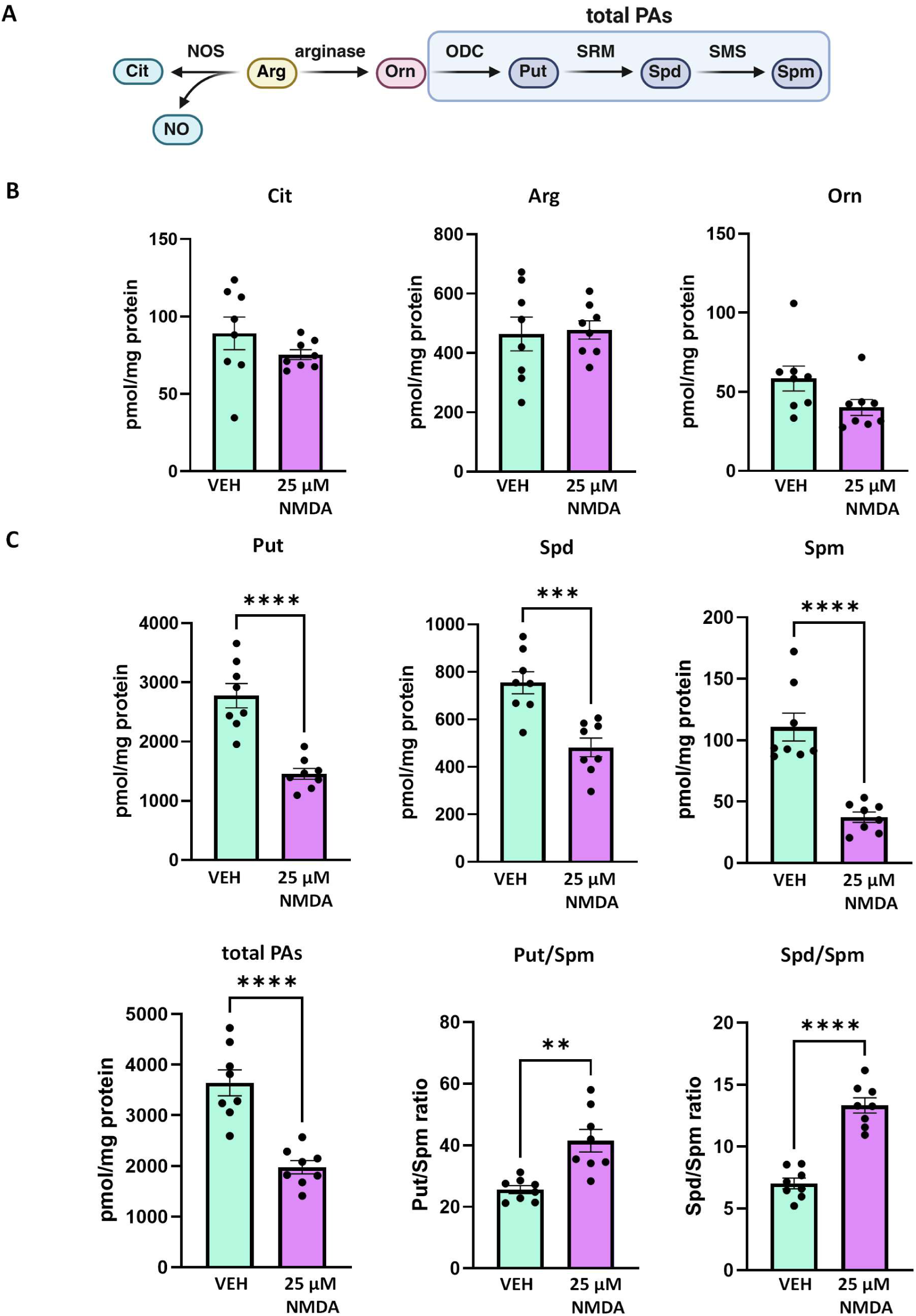
Effect of NMDA treatment on levels of PAs and related AAs in the OHCs. **(A)** Schematic representation of Arg metabolic pathway. **(B)** Measurements of Arg and related AA levels in the VEH- and 25 µM NMDA-exposed OHCs. No statistically significant differences in Cit levels were observed between VEH- and NMDA-exposed OHCs (89.14 ± 10.59 pmol/mg protein vs. 75.32 ± 3.18 pmol/mg protein, respectively; p = 0.2318, n = 8 biological replicates (individual cultures)). No statistically significant differences in Arg levels were observed between VEH- and NMDA-exposed OHCs (464.19 ± 57.14 pmol/mg protein vs. 477.63 ± 30.73 pmol/mg protein, respectively; p = 0.8388, n = 8 biological replicates (individual cultures)). No statistically significant differences in Orn levels were observed between VEH- and NMDA-exposed OHCs (58.47 ± 7.85 pmol/mg protein vs. 40.16 ± 5.08 pmol/mg protein, respectively; p = 0.0703, n = 8 biological replicates (individual cultures)). **(C)** Measurements of PA levels in the VEH- and 25 µM NMDA-exposed OHCs. A statistically significant decrease in Put levels was observed in NMDA-exposed OHCs compared to VEH-exposed OHCs (1455.99 ± 91.89 pmol/mg protein vs. 2777.12 ± 204.15 pmol/mg protein, respectively; p < 0.0001, n = 8 biological replicates (individual cultures)). A statistically significant decrease in Spd levels was observed in NMDA-exposed OHCs compared to VEH-exposed OHCs (482.26 ± 39.32 pmol/mg protein vs. 754.18 ± 46.41 pmol/mg protein, respectively; p = 0.0005, n = 8 biological replicates (individual cultures)). A statistically significant decrease in Spm levels was observed in NMDA-exposed OHCs compared to VEH-exposed OHCs (37.33 ± 4.20 pmol/mg protein vs. 110.79 ± 11.32 pmol/mg protein, respectively; p < 0.0001, n = 8 biological replicates (individual cultures)). A statistically significant decrease in total PAs levels was observed in NMDA-exposed OHCs compared to VEH-exposed OHCs (1975.58 ± 130.11 pmol/mg protein vs. 3642.09 ± 257.12 pmol/mg protein, respectively; p < 0.0001, n = 8 biological replicates (individual cultures)). A statistically significant increase in Put/Spm ratio was observed in NMDA-exposed OHCs compared to VEH-exposed OHCs (41.51 ± 3.65 vs. 25.61 ± 1.28, respectively; p = 0.0011, n = 8 biological replicates (individual cultures)). A statistically significant increase in Spd/Spm ratio was observed in NMDA-exposed OHCs compared to VEH-exposed OHCs (13.32 ± 0.61 vs. 7.025 ± 0.43, respectively; p < 0.0001, n = 8 biological replicates (individual cultures)). Data are presented as mean ± SEM. **p < 0.01, ***p < 0.001, ****p < 0.0001, unpaired t-test. Abbreviations: AA/AAs – amino acid/amino acids; Arg – arginine; Cit – citrulline; NMDA – N-methyl-D-aspartate; NO – nitric oxide; NOS – nitric oxide synthase; ODC – ornithine decarboxylase; OHC/OHCs – organotypic hippocampal slice/slices; Orn – ornithine; PA/Pas – polyamine/polyamines; Put – putrescine; SMS – spermine synthase; Spd – spermidine; Spm – spermine; SRM – spermidine synthase; VEH – vehicle.

The levels of Cit, Arg and or Orn did not differ between vehicle- and NMDA-exposed slices (Fig. 4B). In contrast, NMDA treatment significantly reduced the individual levels of all three PAs and their total content (Fig. 4C). These decreases were accompanied by changes in the ratios of individual PAs (increase in Put/Spm and Spd/Spm (Fig. 4C) and decrease in Put/Spd (Fig. S5A)), and a significant reduction in N^1^-AcSpm, but not N^1^-AcSpmd levels (Fig. S5B).

Together, the selective decreases in PAs, without corresponding changes in related AAs, suggest regulated modulation of PAs homeostasis rather than nonspecific metabolite loss associated with cell death, implicating PA metabolism in the hippocampal response to NMDA- induced excitotoxicity.

### Inhibition of ODC increases NMDA-induced neuronal injury in CA2

Whole-slice measurements reveal neither the regional specificity of metabolic changes nor their contribution to differential vulnerability to NMDA-induced injury. We therefore took advantage of OHCs, which largely preserve hippocampal architecture, cellular interactions, and subregional properties (Stoppini et al., 1991), allowing spatially resolved assessment of cell injury (Skweres et al., 2026). We pharmacologically modulated PAs synthesis in this NMDA excitotoxicity model and assessed damage using the cell injury probe, PI, separately in CA1 and CA2, the subregions with the most contrasting PAs-synthesizing enzyme expression profiles and also the greatest difference in NMDA sensitivity.

We first used DFMO to assess effects of ODC activity inhibition (Pegg et al., 1987) on regional sensitivity to NMDA. NMDA alone (25 μM) induced clearly greater neuronal injury in CA1 than in CA2 (Fig. 5A). Although DFMO alone (0.2 or 1 mM) caused no detectable injury in any subregion, both concentrations visibly increased NMDA-induced injury in CA2 (Fig. 5B). ANOVA of PI intensity in the sp of CA1 and CA2 revealed a significant effect of region (Fig. 5C). Tukey’s multiple-comparisons test confirmed significantly lower PI intensity in CA2 than in CA1 after NMDA exposure. DFMO significantly increased NMDA-induced PI intensity in CA2 at both concentrations but had no significant effect in CA1 (Fig. 5C). The complete results of Tukey’s multiple comparisons test are provided in Table S1.

**Fig. 5.**
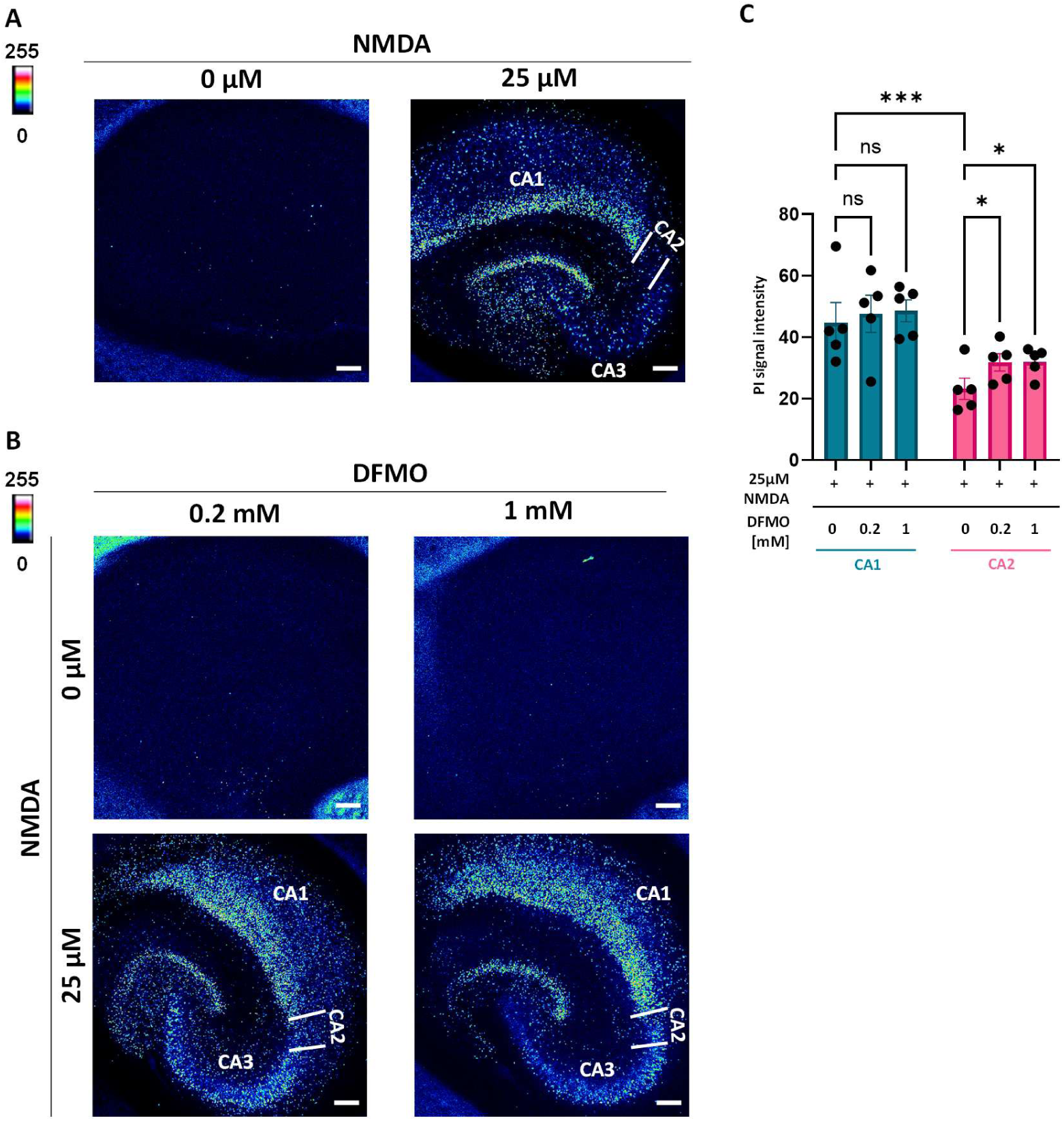
Effect of DFMO exposure on NMDA-induced injury in CA1 and CA2 regions in OHCs. **(A)** Slices were exposed to either VEH (left) or 25 μM NMDA and stained with PI (pseudocolored for intensity ((low) 0-255 (high)). Representative images are shown. Scale bars: 200 μm. **(B)** Slices were exposed to 0.2 mM DFMO (top left), 1 mM DFMO (top right), 25 μM NMDA combined with 0.2 mM DFMO (bottom left) or 25 μM NMDA combined with 1 mM DFMO (bottom right) and stained with PI (pseudocolored for intensity ((low) 0-255 (high)). Representative images are shown. Scale bars: 200 μm. **(C)** Mean PI fluorescence intensity was quantified in the CA1 and CA2 regions of OHCs exposed to 25 μM NMDA alone or in combination with DFMO (0.2 or 1 mM). A two-way mixed ANOVA with region as the repeated-measures factor, revealed a significant main effect of region (F(1, 4) = 40.66, p = 0.0031), but no significant main effect of treatment (F(2, 8) = 1.198; p = 0.3506) or treatment × region interaction (F(2, 8) = 1.919; p = 0.2086). Tukey’s multiple comparison test identified no significant differences in PI intensity between CA1-25 μM NMDA vs. CA1-25 μM NMDA + 0.2 mM DFMO (p = 0.796), no significant differences in PI intensity between CA1-25 μM NMDA vs. CA1-25 μM NMDA + 1 mM DFMO (p = 0.57), significantly higher PI intensity in CA2-25 μM NMDA + 0.2 mM DFMO vs. CA2-25 μM NMDA (p = 0.04), significantly higher PI intensity in CA2-25 μM NMDA + 1 mM DFMO vs. CA2-25 μM NMDA (p = 0.036) and significantly lower PI intensity in CA2-25 μM NMDA vs. CA1-25 μM NMDA (p = 0.0001). The complete results of Tukey’s multiple comparisons test are provided in Table S1. Data are presented as mean ± SEM. n = 5 biological replicates (individual cultures), *p < 0.05, ***p < 0.001, Tukey’s post-hoc test. Abbreviations: DFMO – difluoromethylornithine, inhibitor of ornithine decarboxylase; NMDA – N-methyl-D-aspartate, ns – not significant; OHC/OHCs – organotypic hippocampal culture/cultures; PI – propidium iodide, VEH – vehicle.

Together, these findings suggest that ODC-dependent Put synthesis contributes to CA2 resistance to NMDA-induced excitotoxic injury, consistent with the enrichment of ODC in CA2.

### Inhibition of SMS reduces NMDA-induced neuronal injury in CA2

Next, we examined whether inhibition of SMS, another CA2-enriched enzyme involved in PAs synthesis alters regional vulnerability to NMDA-induced injury in OHCs. After screening several concentrations of the SMS inhibitor, APCHA (Beppu et al., 1995), we selected 1 mM, which caused no visible injury when applied alone (Fig. S6A). OHCs were then treated with APCHA in combination with 25 μM NMDA either alone or with 100 μM exogenous Spm (Fig. 6A).

**Fig. 6.**
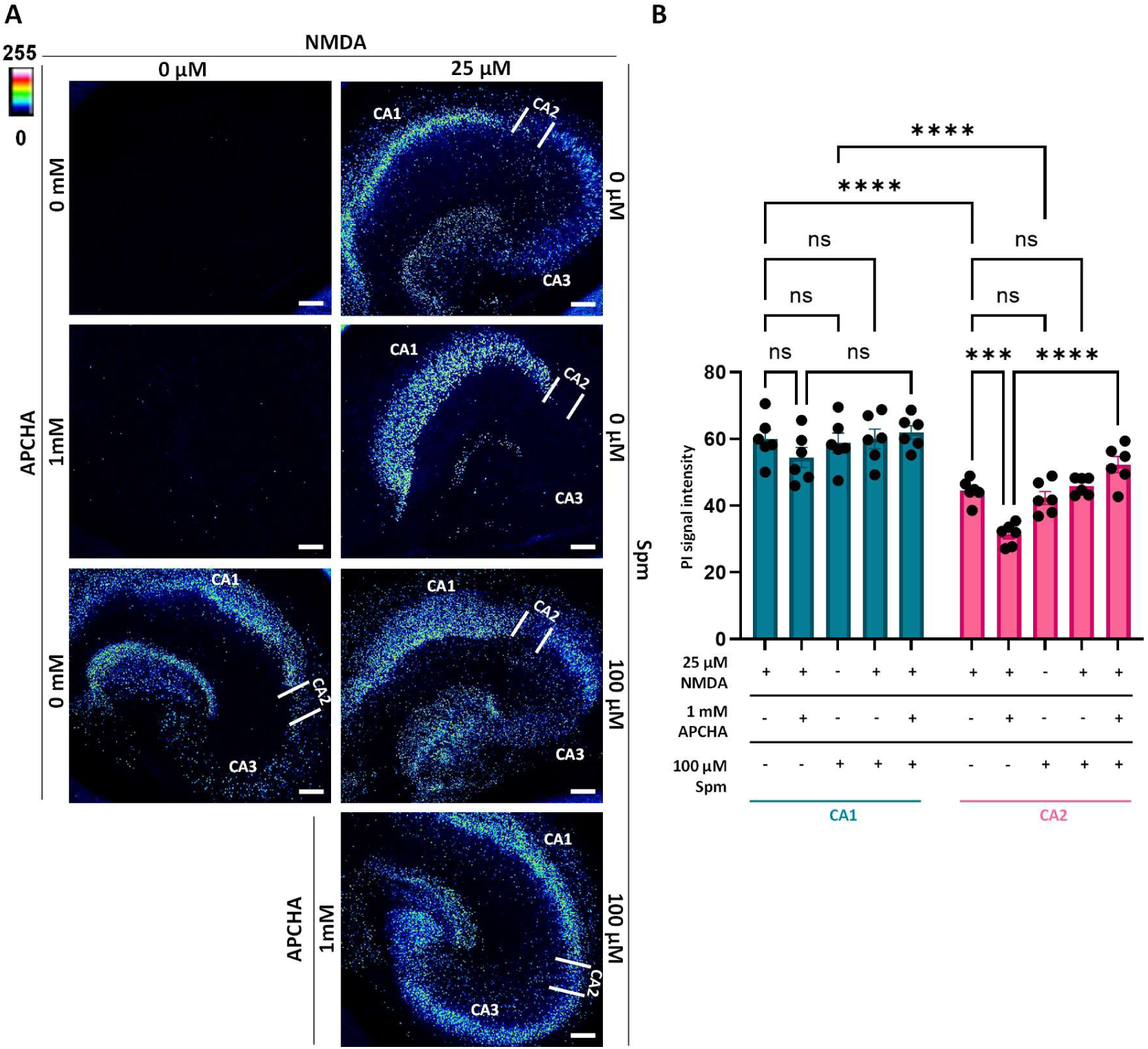
Effect of APCHA and/or Spm exposure on NMDA-induced injury in CA1 and CA2 regions in OHCs. **(A)** Slices were exposed to VEH (top left), 25 μM NMDA (top right), 1 mM APCHA (second row, left), 25 μM NMDA combined with 1 mM APCHA (second row, right), 100 μM Spm (third row, left), 25 μM NMDA combined with 100 μM Spm (third row, right), or 25 μM NMDA combined with 1 mM APCHA and 100 μM Spm (bottom right) and stained with PI (pseudocolored for intensity ((low) 0-255 (high)). Representative images are shown. Scale bars: 200 μm. **(B)** Mean PI fluorescence intensity was quantified in the CA1 and CA2 regions of OHCs exposed to 25 μM NMDA or 100 μM Spm, alone or in the presence or absence of APCHA. A two-way mixed ANOVA with region as the repeated-measures factor, revealed a significant main effect of region (F(1, 5) = 207.9, p < 0.0001), treatment (F(4, 20) = 12.07, p < 0.0001), and region x treatment interaction (F(4, 20) = 4.739, p = 0.0075). Tukey’s multiple comparison test identified no significant differences in PI intensity between CA1-25 μM NMDA vs. CA1-25 μM NMDA + 1 mM APCHA (p = 0.329), no significant differences in PI intensity between CA1-25 μM NMDA vs. CA1-100 μM Spm (p > 0.999), no significant differences in PI intensity between CA1-25 μM NMDA vs. CA1-25 μM NMDA + 100 μM Spm (p > 0.999), no significant differences in PI intensity between CA1-25 μM NMDA + 1 mM APCHA vs. CA1-25 μM NMDA + 1 mM APCHA + 100 μM Spm (p = 0.0672), significantly lower PI intensity in CA2-25 μM NMDA + 1 mM APCHA vs. CA2-25 μM NMDA (p = 0.0003), no significant differences in PI intensity between CA2-25 μM NMDA vs CA2-100 μM Spm (p = 0.99), no significant differences in PI intensity between CA2-25 μM NMDA vs CA2-25 μM NMDA + 100 μM Spm (p = 0.9996), significantly higher PI intensity in CA2-25 μM NMDA + 1 mM APCHA + 100 μM Spm vs. CA2-25 μM NMDA + 1 mM APCHA (p < 0.0001), significantly lower PI intensity in CA2-25 μM NMDA vs. CA1-25 μM NMDA (p < 0.0001) and significantly lower PI intensity in CA2-100 μM Spm vs CA1-100 μM Spm (p < 0.0001). The complete results of Tukey’s multiple comparisons test are provided in Table S2. Data are presented as mean ± SEM. n = 6 biological replicates (individual cultures), ***p < 0.001, ****p < 0.0001, Tukey’s post-hoc test. Abbreviations: APCHA - N^1^-cyclohexylpropane-1,3-diamine, inhibitor of spermine synthase; NMDA – N-methyl-D-aspartate, ns – not significant; OHC/OHCs – organotypic hippocampal culture/cultures; PI – propidium iodide; Spm – spermine, VEH – vehicle.

ANOVA of PI intensity in the CA1 and CA2 sp revealed significant effects of region and treatment and a significant region × treatment interaction (Fig. 6B), indicating differential responses of CA1 and CA2 to treatment. APCHA reduced NMDA-induced injury in CA2 but not CA1 (Fig. 6A, B). Exogenous Spm alone induced injury in both CA1 and CA2, with PI signal intensities comparable to those induced by NMDA within each region, but combining Spm with NMDA did not significantly potentiate injury relative to NMDA alone (Fig. 6A,B). Spm, although attenuated APCHA-mediated protection against NMDA-induced injury in CA2 (Fig. 6A), significantly increasing PI intensity relative to NMDA + APCHA in this region, with no significant effect on PI intensity in CA1 (Fig. 6B). The complete results of Tukey’s multiple comparisons test are provided in Table S2.

Together, these findings suggest that SMS-dependent Spm synthesis contributes to the residual sensitivity of CA2 neurons to NMDA-induced injury, with SMS inhibition further enhancing their resistance. The attenuation of APCHA-mediated protection in CA2 by exogenous Spm further supports this interpretation, as does the absence of an APCHA effect on NMDA-induced injury in CA1, where SMS expression is limited.

### Blocking of GluN2B-containing NMDARs attenuates NMDA-induced neuronal injury, but exogenous Spm abolishes this protective effect

Given the attenuation of APCHA-mediated protection against NMDA-induced injury in CA2 by exogenous Spm, we next examined whether this effect involves Spm-mediated potentiation of NMDAR activity. Spm can act as a positive allosteric modulator of GluN2B-containing NMDARs (Mony et al., 2011). We therefore treated OHCs with NMDA and/or Spm in the presence or absence of ifenprodil, a selective, noncompetitive antagonist of GluN2B-containing NMDARs (Perin-Dureau et al., 2002). After screening several concentrations of ifenprodil, we selected 15 μM, which caused no visible injury when applied alone (Fig. S6B). OHCs were then treated with ifenprodil in combination with 25 μM NMDA and/or 100 μM exogenous Spm (Fig. 7A).

**Fig. 7.**
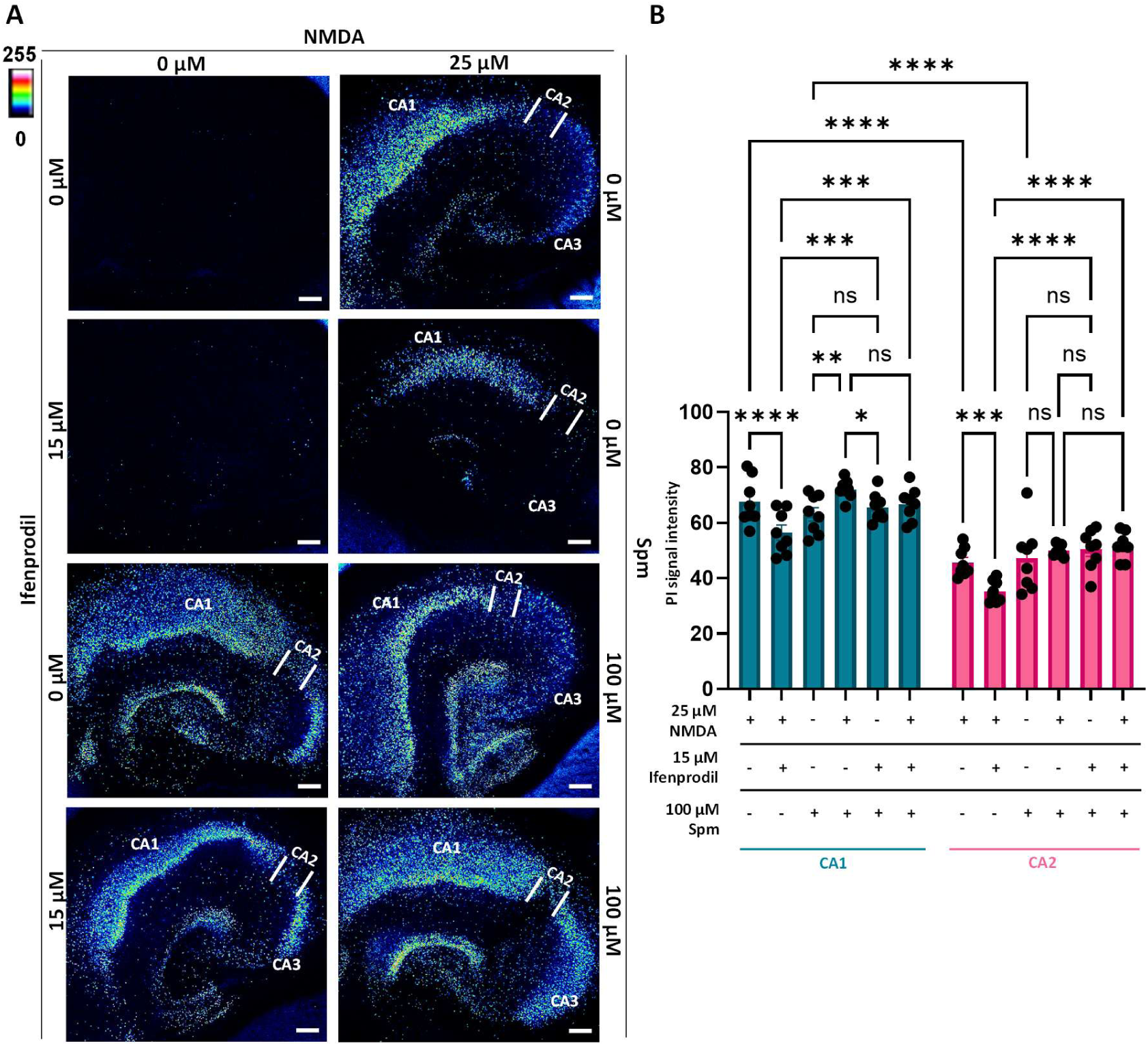
Effect of ifenprodil and/or Spm exposure on NMDA-induced injury in CA1 and CA2 regions in OHCs. **(A)** Slices were exposed to VEH (top left), 25 μM NMDA (top right), 15 μM ifenprodil (second row, left), 25 μM NMDA combined with 15 μM ifenprodil (second row, right), 100 μM Spm (third row, left), 25 μM NMDA combined with 100 μM Spm (third row, right), 15 μM ifenprodil combined with 100 μM Spm (bottom, right), or 25 μM NMDA combined with 15 μM ifenprodil and 100 μM Spm (bottom right) and stained with PI (pseudocolored for intensity ((low) 0-255 (high)). Representative images are shown. Scale bars: 200 μm. **(B)** Mean PI fluorescence intensity was quantified in the CA1 and CA2 regions of OHCs exposed to 25 μM NMDA or 100 μM Spm, alone or in the presence or absence of ifenprodil. A two-way mixed ANOVA with region as the repeated-measures factor, revealed a significant main effect of region (F(1, 7) = 247.7, p < 0.0001), treatment (F(5, 35) = 9.872, p < 0.0001), and region x treatment interaction (F(5, 35) = 3.525, p = 0.011). Tukey’s multiple comparison test identified significantly lower PI intensity in CA1-25 μM NMDA + 15 μM ifenprodil vs. CA1-25 μM NMDA (p < 0.0001), significantly higher PI intensity in CA1-15 μM ifenprodil + 100 μM Spm vs CA1-25 μM NMDA + 15 μM ifenprodil (p = 0.0009), significantly higher PI intensity in CA1-25 μM NMDA + 15 μM ifenprodil + 100 μM Spm vs CA1-25 μM NMDA + 15 μM ifenprodil (p = 0.0001), significantly higher PI intensity in CA1-25 μM NMDA + 100 μM Spm vs CA1-100 μM Spm (p = 0.001), no significant differences in PI intensity between CA1-100 μM Spm vs CA1-15 μM ifenprodil + 100 μM Spm (p = 0.951), significantly lower PI intensity in CA1-15 μM ifenprodil + 100 μM Spm vs CA1-25 μM NMDA + 100 μM Spm (p = 0.0472), no significant differences in PI intensity between CA1-25 μM NMDA + 100 μM Spm vs CA1-25 μM NMDA + 15 μM ifenprodil + 100 μM Spm (p = 0.1768), significantly lower PI intensity in CA2-25 μM NMDA + 15 μM ifenprodil vs CA2-25 μM NMDA (p = 0.0001), significantly higher PI intensity in CA2-15 μM ifenprodil + 100 μM Spm vs CA2-25 μM NMDA + 15 μM ifenprodil (p < 0.0001), significantly higher PI intensity in CA2-25 μM NMDA + 15 μM ifenprodil + 100 μM Spm vs CA2-25 μM NMDA + 15 μM ifenprodil (p < 0.0001), no significant differences in PI intensity between CA2-100 μM Spm vs CA2-25 μM NMDA + 100 μM Spm (p = 0.9267), no significant differences in PI intensity between CA2-100 μM Spm vs CA2-15 μM ifenprodil + 100 μM Spm (p = 0.8392), no significant differences in PI intensity between CA2-25 μM NMDA + 100 μM Spm vs CA2-15 μM ifenprodil + 100 μM Spm (p > 0.999), no significant differences in PI intensity between CA2-25 μM NMDA + 100 μM Spm vs CA2-25 μM NMDA + 15 μM ifenprodil + 100 μM Spm (p = 0.9988), significantly lower PI intensity in CA2-25 μM NMDA vs CA1-25 μM NMDA (p < 0.0001), significantly lower PI intensity in CA2-100 μM Spm vs CA1-100 μM Spm (p < 0.0001). The complete results of Tukey’s multiple comparisons test are provided in Table S3. Data are presented as mean ± SEM. n = 6 biological replicates (individual cultures), *p < 0.05, **p < 0.01, ***p < 0.001, ****p < 0.0001, Tukey’s post-hoc test. Abbreviations: NMDA – N-methyl-D-aspartate, ns – not significant; OHC/OHCs – organotypic hippocampal culture/cultures; PI – propidium iodide; Spm – spermine, VEH – vehicle.

ANOVA of PI intensity in the CA1 and CA2 revealed significant effects of region and treatment and a significant region × treatment interaction (Fig. 7B), indicating differential responses of CA1 and CA2 to treatment. Although ifenprodil reduced NMDA-induced injury in both regions (Fig. 7A), as indicated by significantly lower PI signal intensities in NMDA + ifenprodil-treated slices than in slices exposed to NMDA alone (Fig. 7B), it did not alter injury associated with Spm exposure, either alone or in combination with NMDA, in either CA1 or CA2 (Fig. 7A). Specifically, no significant differences in PI intensity were observed between Spm and Spm + ifenprodil or between Spm + NMDA and Spm + NMDA + ifenprodil in either region (Fig. 7B). In addition, PI fluorescence intensities were significantly higher following Spm + ifenprodil treatment than following NMDA + ifenprodil treatment in both CA1 and CA2 (Fig. 7B), showing that, unlike NMDA-induced toxicity, toxicity induced by Spm alone does not involve a substantial GluN2B-dependent excitotoxic component. Moreover, PI intensities following NMDA + ifenprodil + Spm treatment were significantly higher than those following NMDA + ifenprodil treatment in both CA1 and CA2 (Fig. 7A,B), indicating attenuation of ifenprodil-mediated protection in the presence of exogenous Spm. Importantly, NMDA potentiated Spm toxicity only in CA1, where PI fluorescence intensity was significantly higher following Spm + NMDA treatment than following Spm or Spm + ifenprodil treatment (Fig. 7A,B). This suggests a region-specific interaction between exogenous Spm and NMDA, consistent with the enrichment of GluN2B in CA1 pyramidal neurons and the previously reported increase in sensitivity to NMDA in the presence of Spm (Butler et al., 2010). Notably, the difference between Spm alone and Spm + NMDA in CA1 reached statistical significance in the ifenprodil experimental series but not in the APCHA experimental series (Fig. 6B), possibly reflecting the longer Spm exposure in the former experimental setup (4 h vs 3 h, respectively). The complete results of Tukey’s multiple comparisons test are provided in Table S3.

Together, these findings support a contribution of GluN2B-containing NMDARs to NMDA-induced injury in both regions. However, ifenprodil did not significantly reduce injury induced by NMDA in combination with Spm in CA2. Therefore, these experiments do not support a model where GluN2B-mediated potentiation is the mechanism underlying the attenuation of APCHA-mediated protection in CA2 by exogenous Spm.

Collectively, our findings identify a novel metabolic phenotype of CA2 pyramidal neurons, with selective ARG2, enriched ODC and SMS, and low nNOS expression. ARG2 loss reduces hippocampal PA levels, implicating CA2 in their homeostasis. In rat OHCs, NMDA decreases PA levels without altering related AAs, while ODC and SMS inhibition, respectively increase and decrease NMDA-induced injury selectively in CA2. Exogenous Spm damages both CA1 and CA2 and attenuates APCHA-mediated protection in CA2, supporting a contribution of SMS-dependent Spm synthesis to its residual NMDA sensitivity. Ifenprodil reduced NMDA-induced injury, but Spm abolished this protection, suggesting that Spm counteracts SMS inhibition-mediated CA2 neuroprotection primarily through mechanisms other than GluN2B potentiation. Together, these findings reveal the distinctive organization of PA metabolism in CA2 and implicate it in the functional phenotype of this region’s pyramidal neurons, particularly their unique response to excitotoxic injury.

## DISCUSSION

### A distinctive metabolic phenotype of CA2 pyramidal neurons

We identified a previously unrecognized metabolic phenotype of hippocampal CA2 pyramidal neurons: substantial enrichment in enzymes of the Arg-to-PAs synthesis pathway, suggesting a particularly high capacity for Arg-derived PA synthesis in these cells. Low expression of nNOS a potential competitor for Arg provides further supports this interpretation. ARG2, which catalyzes the first step of this pathway, and whose expression was restricted to CA2, appears to drive this metabolic specialization. To our knowledge, such strict CA2 specificity has not been reported previously, although Arg2 was identified among the transcripts distinguishing CA2 from other hippocampal regions in mice (Cembrowski et al., 2016; Farris et al., 2019) and rats (Machado et al., 2022). Also, Polis and colleagues detected ARG2 protein in CA2 pyramidal neurons by immunohistochemistry in the 3xTg mouse model of Alzheimer’s disease, but, unexpectedly, reported no detectable ARG2 in pyramidal neurons in non-transgenic mice (Polis et al., 2018).

The functional importance of this pathway was supported by the disproportional effect of ARG2 loss on hippocampal PA content: despite ARG2 expression being restricted to CA2 pyramidal neurons, its loss reduced total PA content in whole-hippocampus homogenates by ∼35%, far exceeding CA2 relative size (Attili et al., 2019). Also, the distinct response of striatal medium spiny neurons to ARG2 loss (Nalepa et al., 2026), together with brain’s dependence on local PA synthesis rather than uptake from the circulation (Shin et al., 1985; Weiss et al., 2023) support CA2-specific mechanism. This presumption is further supported by the report of Laube and colleagues, who studied Spd/Spm-like immunoreactivity throughout the rat brain and identified a strong, distinctive signal in the CA2 pyramidal layer (Laube et al., 2002). This biochemical specialization suggests that PAs support particular metabolic requirements of CA2 pyramidal neurons and contribute to their functional differences from neighbouring regions. Consistent with this interpretation, we show that inhibition of PA synthesis enzymes modulates NMDA sensitivity in CA2 but not in CA1, the CA region most distinct from CA2 (Farris et al., 2019).

The effect of ARG2 loss on hippocampal PAs different clearly from that in the striatum, where the PA balance shifted towards higher PAs without changes in total PA content (Nalepa et al., 2026). Together with absent or negligible ARG2 expression in several other examined regions (Bichell et al., 2017; Nalepa et al., 2025), this suggests that neuronal ARG2 supports the metabolic specialization of selected neuronal populations, with its effects depending on the local organization of PA homeostasis. Despite these differences, hippocampal Orn levels remained unchanged, similarly as in the striatum (Nalepa et al., 2026), whole brain (Xu et al., 2023), plasma (Didelija et al., 2017; Shi et al., 2001) and peripheral tissues, including intestine (Marini et al., 2011) of Arg2^-/-^ mice. This suggests that ARG2 loss does not necessarily result in altered Orn abundance, possibly because compensatory pathways that are able to maintain Orn availability, although they fail to prevent PA dyshomeostasis, as demonstrated in the striatum (Nalepa et al., 2026).

### Regional organization of PAs synthesis

The differing distributions of individual enzymes suggest that the Arg-to-Spm pathway is not present as a complete enzymatic sequence throughout the hippocampal pyramidal neuron subpopulations. While CA2 expresses the full enzymatic machinery at high levels, indicating substantial PA biosynthetic potential, neighbouring regions, particularly CA1, exhibit limited potential, suggesting distinct requirements for individual PAs, and/or greater dependence on PA uptake. However, the mechanisms of PA uptake, particularly into neurons, remain incompletely understood. The prevailing view is that PAs are synthesized mainly in neurons (Kilpeläinen et al., 2000; Krauss et al., 2006, 2007), but accumulate predominantly in astrocytes (Biedermann et al., 1998; Laube et al., 1997; Skatchkov et al., 2000), implicating neuronal release followed by astrocytic uptake (Rieck et al., 2022). CA2 appears to represent an exception, as the distinctive Spd/Spm-like immunoreactivity in its pyramidal layer suggests preferential accumulation of these PAs in its neurons (Laube et al., 2002).

In contrast to the prevailing view of astrocytic accumulation, a recent study using the bioorthogonal Put probe iPUT demonstrated preferential neuronal labelling in neuron– astrocyte co-cultures and marked accumulation in pyramidal neuron somata in CA1-CA3 subregions in acute mouse hippocampal slices (Owczarek et al., 2026). This may point to a distinct cellular distribution of Put compared with the better studied Spd and Spm. Given the brain’s reliance on local PA synthesis (Weiss et al., 2023), this capacity for neuronal uptake may indicate that CA2, where ARG2 and ODC are highly enriched, supplies Put to neighbouring region’s neurons for Spd synthesis. Consistent with this possibility, we found high SRM expression in these neurons, in contrast to the upstream enzymes. Although, despite CA1 and CA3 pyramidal neurons appear capable of synthesizing Spd, they show little evidence of its accumulation in contrast to CA2 pyramidal cells (Laube et al., 2002).

### NMDA-induced remodelling of PA homeostasis

NMDA exposure reduced all three PAs in OHCs without detectable changes in Cit, Arg or Orn, suggesting controlled modulation of PA homeostasis rather than nonspecific metabolite loss from damaged cells. Changes in PA ratios further indicated that depletion was not uniform across individual PAs. Given the opposing effects of ODC- and SMS-directed inhibition on the CA2 response to NMDA, this shift may reflect an adaptive response involving not only PA depletion but also a shift towards potentially survival-promoting PA ratios – higher Put/Spm and Spd/Spm.

Unlike reports of increased Put and/or ODC induction following NMDAR activation or excitotoxic injury *in vivo* (Camón et al., 2001; Giménez-Llort et al., 1997; Mácha et al., 2024; Porcella et al., 1991) and *in vitro* (Lombardi et al., 1993; Siddiqui & Iqbal, 1994), NMDA reduced all three PAs in our study. This finding partially corresponds to the early response to intrastriatal injection of NMDA, where Spd and Spm depletion preceded ODC induction (Porcella et al., 1991) in the striatum, another ARG2-enriched brain region (Bichell et al., 2017; Nalepa et al., 2025). NMDA was also shown to induce striatal Spd and Spm release without affecting extracellular Put (Porcella et al., 1992), suggesting that PA export may contribute to this depletion. Although whole-slice measurements do not represent region-specific alterations, the enrichment of PA-related features in CA2 suggests that this region may contribute disproportionately to the observed PA changes.

### Opposing effects of targeting distinct steps in the PA synthesis pathway

DFMO and APCHA modulated the response to NMDA in CA2 but not CA1 consistent with our finding that their respective target enzymes, ODC and SMS, are enriched in CA2, but absent or barely detectable in CA1 pyramidal neurons. Their opposing effects in CA2 highlight the distinct contributions of individual PAs to excitotoxic vulnerability: local synthesis of Put and possibly Spd appears to support CA2 resistance, while Spm production may contribute to its residual vulnerability in our model. These findings further support the possibility that NMDA-induced changes in PA ratios, particularly the most pronounced Spm reduction, reflect an adaptive response aimed at limiting excitotoxic damage.

Previous studies have reported differential effects of ODC modulation on hippocampal neuronal survival. DFMO protected CA1 neurons in ischemic gerbils (Kindy et al., 1994) and an ethanol withdrawal OHC model, but did not alter direct NMDA-induced CA1 injury in the latter study (Gibson et al., 2003). In contrast, DFMO exacerbated CA3 neuronal damage in a rat model of transient forebrain ischemia (Zoli et al., 1993), while ODC-overexpressing mice exhibited an elevated seizure threshold and greater protection against ischemic injury (Halonen et al., 1993). Our study provide a complementary perspective by revealing regional enzymatic organization of hippocampal PA synthesis, raising the question of how local production capacity influences neuronal vulnerability. This heterogeneity indicates that the effects of PA synthesis cannot be considered as uniformly protective or detrimental, and its effects depend on the subregion, insult, and state of PA homeostasis.

DFMO affected CA2 pyramidal neuron vulnerability to NMDA despite extracellular Put availability from B27, a supplement commonly used in OHC and primary neuron culturing. This suggests that exogenous Put does not fully compensate for inhibition of endogenous Put synthesis in CA2. Consistent with this interpretation, iPUT experiments in acute mouse slices demonstrated efficient uptake by hippocampal pyramidal neurons, that was reduced by exposure to NMDA in CA1 but not CA2 (Owczarek et al., 2026). Together, these findings suggest that extracellular Put alone is insufficient to protect CA2 pyramidal neurons against NMDA-induced injury, and endogenous Put synthesis is required for this protective effect.

The present findings should be interpreted in light of the distinct importance of intracellular and extracellular PAs. Extracellular Spm and Spd potentiate NMDAR currents (Araneda et al., 1993) through GluN2B-selective allosteric modulation (Mony et al., 2011), and can exacerbate excitotoxic injury *in vivo* (Munir et al., 1993) and *in vitro* (De Vera et al., 2008; Lombardi et al., 1993; Sparapani et al., 1997), including OHCs, where Spd combined with otherwise non-toxic NMDA concentrations induced CA1 injury (Butler et al., 2010). Our findings, however, suggest that the contribution of Spm to CA2 vulnerability under excitotoxic NMDA exposure may involve intracellular mechanisms. Exogenous Spm attenuated APCHA-induced protection from NMDA-induced excitotoxicity but did not exacerbate injury induced by NMDA alone, consistent with replenishment of the intracellular Spm pool following SMS inhibition rather than enhancement of NMDAR signalling. In addition, exogenous Spm fully neutralized protection by ifenprodil, a GluN2B-selective negative allosteric modulator, which acts at distinct site than Spm (Han et al., 2008). This further supports the involvement of intracellular Spm mechanisms and the importance of endogenous Spm synthesis in CA2 neuronal response to excitotoxicity.

Intracellular Spm can also regulate NMDARs by reducing its opening probability independently of subunit composition (Turecek et al., 2004). Given the neuron’s limited access to circulating PAs and reliance on local synthesis and storage, the enrichment of these processes in hippocampal CA2 pyramidal neurons suggests that this inhibitory mechanism may be particularly prominent in this region. However, APCHA reduced rather than increased CA2 vulnerability to NMDA, suggesting that any NMDAR inhibition by endogenous Spm is masked by other mechanisms through which local Spm synthesis promotes excitotoxic injury.

In summary, we identify a previously undescribed metabolic phenotype of CA2 pyramidal neurons, characterized by a distinct pattern of Arg metabolism-related enzymes indicating substantially enhanced PA biosynthetic capacity, and demonstrate that endogenous PA homeostasis influences their vulnerability to NMDA-induced excitotoxicity. This phenotype distinguishes CA2 from neighbouring subregions, particularly CA1, which is highly sensitive to NMDA, expresses much lower levels of the PA-related proteins, and demonstrates no changes in NMDA-induced injury following modulation of PA synthesis.

### Limitations of the study

Some limitations should be considered when interpreting these findings. PAs were quantified in whole hippocampal or OHC homogenates, without revealing differences between individual subregions and intracellular and extracellular pools. Enzyme enrichment indicates biosynthetic capacity but does not establish metabolic flux or PA transfer between subregions. Thus, although hippocampal ARG2 expression was restricted to CA2 pyramidal neurons and most other enzymes were enriched there, PA reductions following ARG2 loss or NMDA exposure cannot be attributed exclusively to CA2 depletion. Also, the DFMO and APCHA findings require genetic validation and direct PA measurements, preferably in precisely dissected CA1 and CA2, to establish region-specific contributions of individual PAs. Medium-derived Put may partially mask the importance of CA2 synthesis of Put and its potential redistribution to other regions. Finally, the relevance of the functional findings obtained in NMDA-exposed OHCs to adult hippocampal physiology and neuronal injury *in vivo* remains to be established.

## ACKNOWLEDGEMENTS

This work was supported by National Science Centre, Poland (grants number 2023/49/N/NZ4/02660 (AS), 2023/51/B/NZ4/02605 (MW), 2018/30/E/NZ1/00144 (MW)) and by MMRI PAS (Internal Research Fund no FBW-024 (AS), FBW-019 (MW)).

The authors wish to thank Dr Hanna Kozłowska of the Laboratory of Advanced Microscopy Techniques, MMRI PAS, for her assistance with confocal imaging.

## AUTHOR CONTRIBUTIONS

**AS**: Conceptualization; Formal analysis; Funding acquisition; Investigation; Methodology; Project administration; Validation; Visualization; Writing – original draft; Writing – review & editing. **AO**: Investigation, Methodology, Visualization. **MBH**: Investigation, Methodology. **ADuchnowska**: Investigation. **MN**: Investigation, Writing – review & editing. **ADalka**: Investigation. **MR**: Investigation. **MWypchło**: Investigation. **FB**: Investigation. **CC**: Investigation. **ABB**: Conceptualization, Writing – review & editing. **GV**: Supervision. **BZ**: Resources, Supervision, Writing – review & editing. **MWęgrzynowicz**: Conceptualization; Funding acquisition; Methodology; Project administration; Resources; Supervision; Writing – original draft; Writing – review & editing.

## DECLARATION OF GENERATIVE AI AND AI-ASSISTED TECHNOLOGIES IN THE WRITING PROCESS

During the preparation of this work, the authors used Perplexity and OpenAI’s ChatGPT to improve the language and readability of the manuscript. After using these tools, the authors reviewed and edited the content as needed and take full responsibility for the content of the published article.

## ABBREVIATIONS

AA/AAs: amino acid/amino acids
APCHA: N^1^-cyclohexylpropane-1,3-diamine
Arg: arginine
ARG1: arginase 1
ARG2: arginase 2
Arg2^-/-^: arginase 2 knockout mouse line
Arg2^+/+^: arginase 2-positive mouse line
Cit: citrulline
DFMO: difluoromethylornithine
DIV: day in vitro
GluN2B: glutamate n-methyl-d-aspartate receptor subtype 2B
IF: immunofluorescence
IHC: immunohistochemistry
MMRI PAS: Mossakowski Medical Research Institute, Polish Academy of Sciences
N^1^-AcSpd: N^1^-acetyl-spermidine
N^1^-AcSpm: N^1^-acetyl-spermine
NHS: normal horse serum
NMDA: n-methyl-d-aspartate
NMDAR/NMDARs: n-methyl-d-aspartate receptor/receptors
nNOS: neuronal nitric oxide synthase
NO: nitric oxide
NOS: nitric oxide synthase
NOx: nitrates and nitrites
ODC: ornithine decarboxylase
OHC/OHCs: organotypic hippocampal culture/cultures
Orn: ornithine
PI: propidium iodide
PA/PAs: polyamine/polyamines
PBST: phosphate-buffered saline with Tween-20
PFA: paraformaldehyde
Put: putrescine
RGS-14: Regulator of G-protein Signaling 14
slm: stratum lacunosum-moleculare
SMS: spermine synthase
so: stratum oriens
sp: stratum pyramidale
Spd: spermidine
Spm: spermine
sr: stratum radiatum
SRM: spermidine synthase

