## Supplementary data for "Distinct polyamine metabolic signatures underlie differential sensitivity of hippocampal CA1 and CA2 regions to NMDA-induced excitotoxicity"

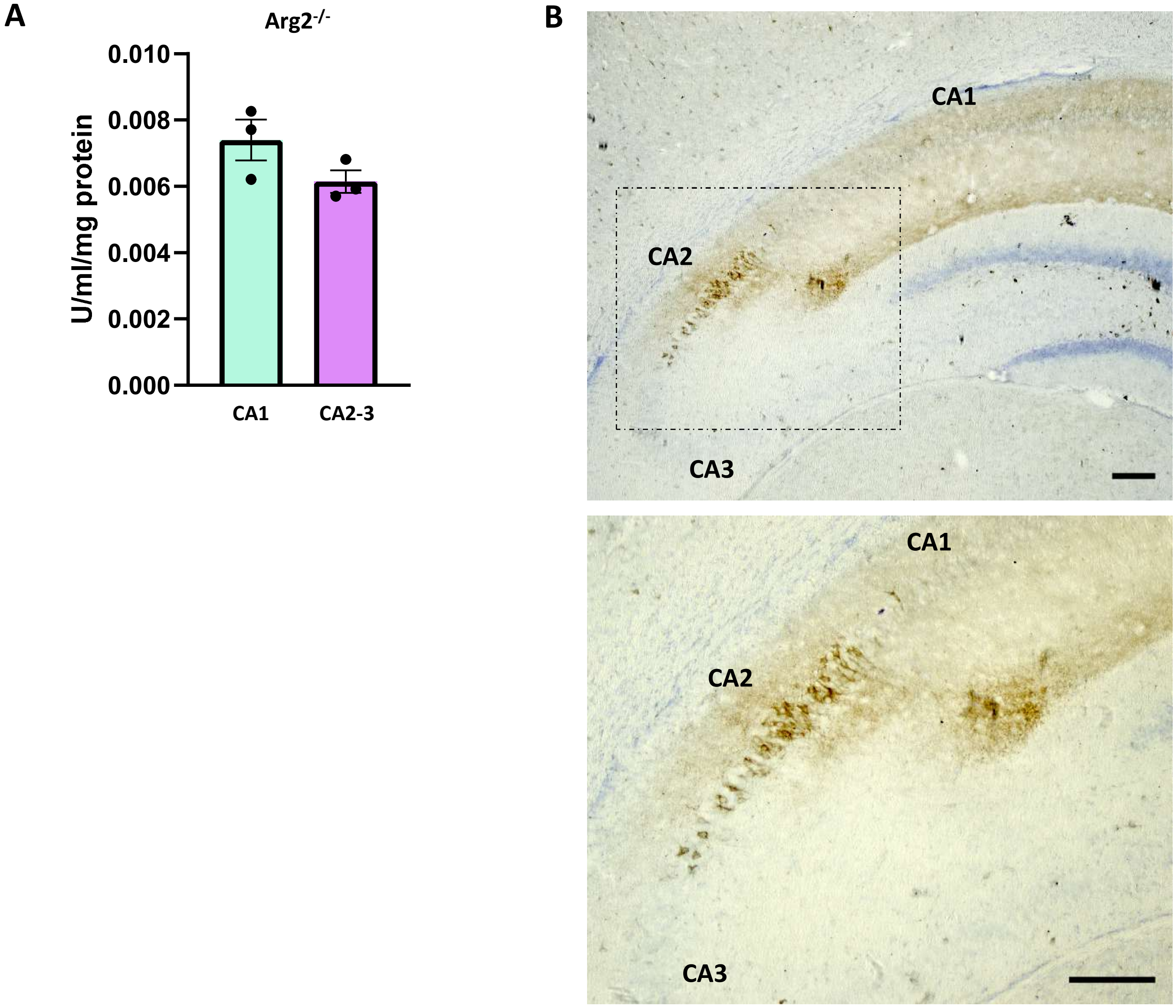


**Fig. S1. Arginase in the hippocampus.** **(A)** Arginase catalytic activity was measured in homogenates of CA2-3 and CA1 hippocampal regions of Arg2$^{-/-}$ mice. No statistically significant differences were observed between CA2-3 and CA1 (0.0061 ± 0.0003 units/ml/mg of protein vs. 0.0074 ± 0.0006, respectively); $p = 0.1481$; $n = 3$ biological replicates (individual animals)). Data are presented as mean ± SEM; $^{**}p < 0.01$, unpaired t-test. **(B)** ARG2 immunohistochemistry in the hippocampus of wild-type rats reveals strong enrichment in the CA2 region. ARG2 localizes to the cell bodies of CA2 pyramidal neurons and their processes, particularly to the stratum lacunosum-moleculare (top panel; enlargement of the corresponding area outlined by a dashed line in the bottom panel). Other hippocampal regions show little to no detectable ARG2 staining. Representative images are shown. Scale bars: 50 µm (top and bottom panels). Abbreviations: ARG2 – arginase 2; Arg2$^{-/-}$ – arginase 2 knockout mouse line.

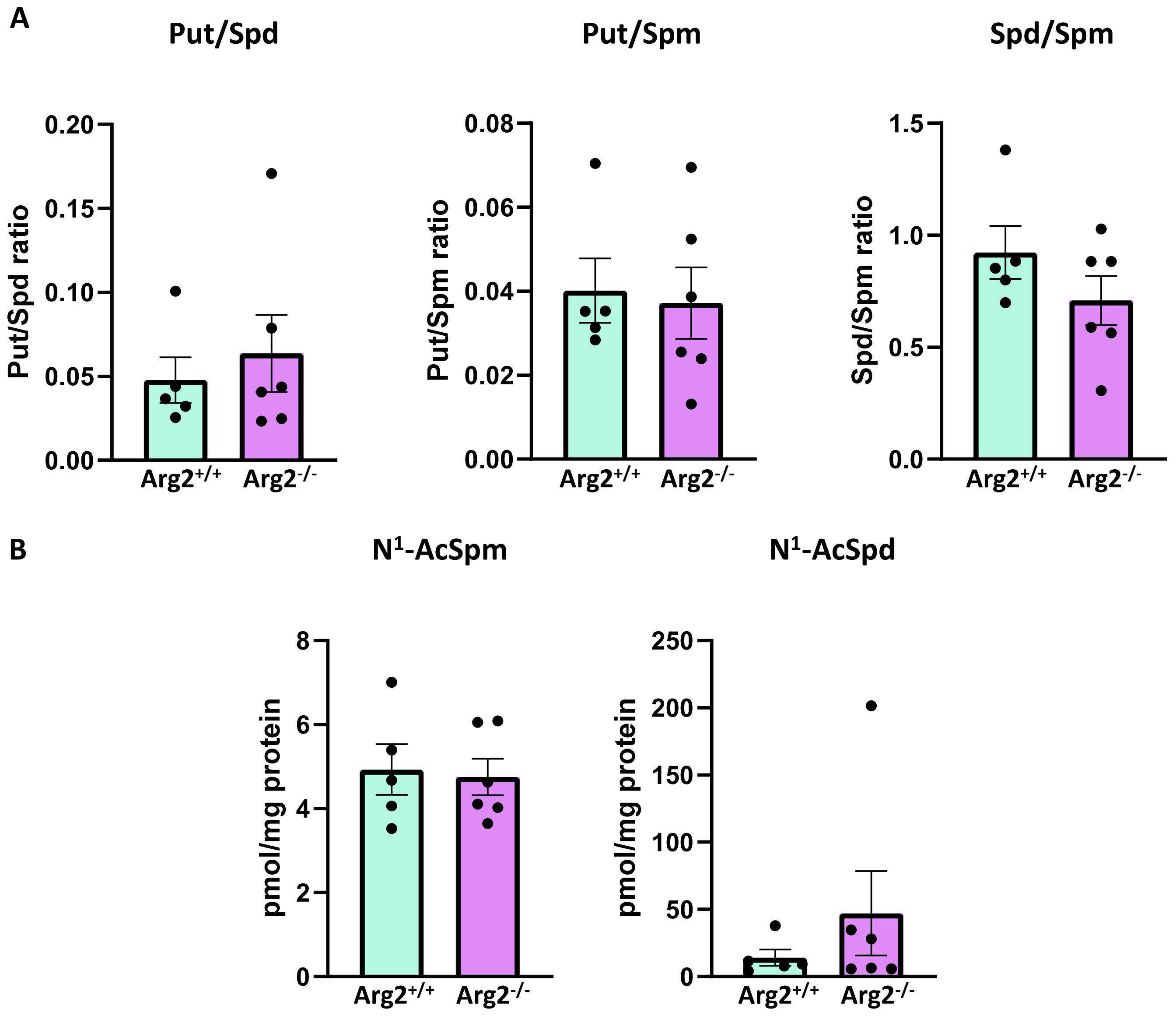


**Fig. S2. Effect of ARG2 loss on PA ratios and levels of PA degradation products in the mouse hippocampus. (A)** Measurements of PA ratios in the hippocampi of $Arg2^{+/+}$ and $Arg2^{-/-}$ mice. No statistically significant differences in Put/Spd ratio were observed between $Arg2^{+/+}$ and $Arg2^{-/-}$ mice (0.0478 ± 0.0136 vs. 0.0637 ± 0.0226, respectively; $p = 0.587$; n = 5–6 biological replicates (individual animals)). No statistically significant differences in Put/Spm ratio were observed between $Arg2^{+/+}$ and $Arg2^{-/-}$ mice (0.0402 ± 0.0076 vs. 0.0372 ± 0.0084, respectively; $p = 0.807$; n = 5–6 biological replicates (individual animals)). No statistically significant differences in Spd/Spm ratio were observed between $Arg2^{+/+}$ and $Arg2^{-/-}$ mice (0.9237 ± 0.1172 vs. 0.7090 ± 0.1099, respectively; $p = 0.216$; n = 5–6 biological replicates (individual animals)). **(B)** Measurements of PA degradation products in the hippocampi of $Arg2^{+/+}$ and $Arg2^{-/-}$ mice. No statistically significant differences in N1-AcSpm levels were observed between $Arg2^{+/+}$ and $Arg2^{-/-}$ mice (4.933 ± 0.623 pmol/mg protein vs. 4.758 ± 0.443 pmol/mg protein, respectively; $p = 0.8156$; n = 5–6 biological replicates (individual animals)). No statistically significant differences in N1-AcSpd levels were observed between $Arg2^{+/+}$ and $Arg2^{-/-}$ mice (14.11 ± 6.00 pmol/mg protein vs. 47.03 ± 31.42 pmol/mg protein, respectively; $p = 0.3724$; n = 5–6 biological replicates (individual animals)). Data are presented as mean ± SEM. Statistical analysis was performed using an unpaired t-test. Abbreviations: $Arg2^{-/-}$ – arginase 2 knockout mouse line; $Arg2^{+/+}$ – arginase 2-positive mouse line; $N^1$-AcSpd – $N^1$-acetyl-spermidine; $N^1$-AcSpm – $N^1$-acetyl-spermine; PA – polyamines; Put – putrescine; Spd – spermidine; Spm – spermine.

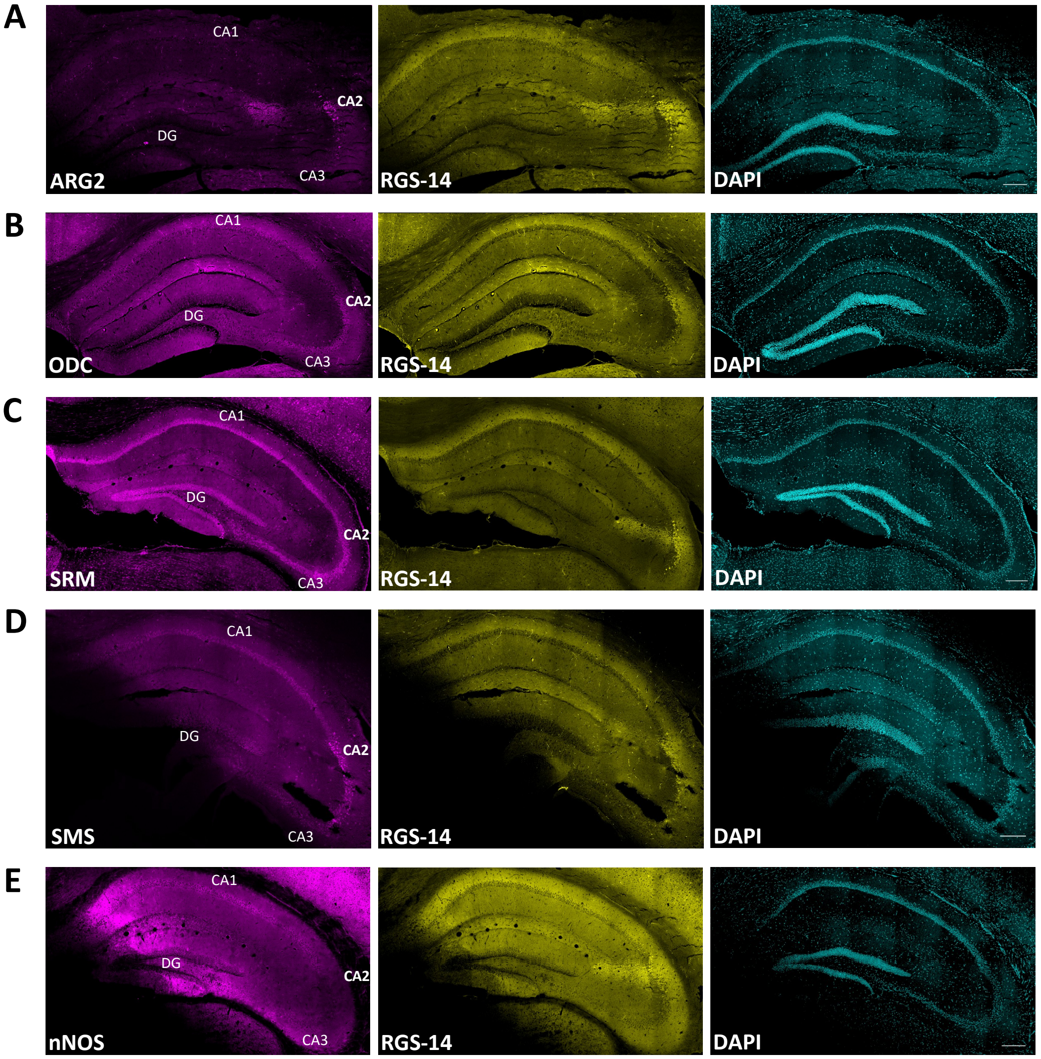


**Fig. S3. Distribution of PA synthesis-related enzymes in the mouse hippocampus (split channels).** Double immunofluorescence staining of mouse hippocampal sections showing **(A)** ARG2, **(B)** ODC, **(C)** SRM, **(D)** SMS **(E)**, and **(E)** nNOS (magenta). Sections were immunostained for the CA2 marker, RGS-14 (yellow) and counterstained with DAPI (cyan). Panels show overview images of the entire hippocampus in separate fluorescence channels. Representative images are shown. Scale bars: 200 µm. Abbreviations: ARG2 – arginase 2; nNOS – neuronal nitric oxide synthase; ODC – ornithine decarboxylase; RGS-14 – Regulator of G-protein Signaling 14; SMS – spermine synthase; SRM – spermidine synthase.

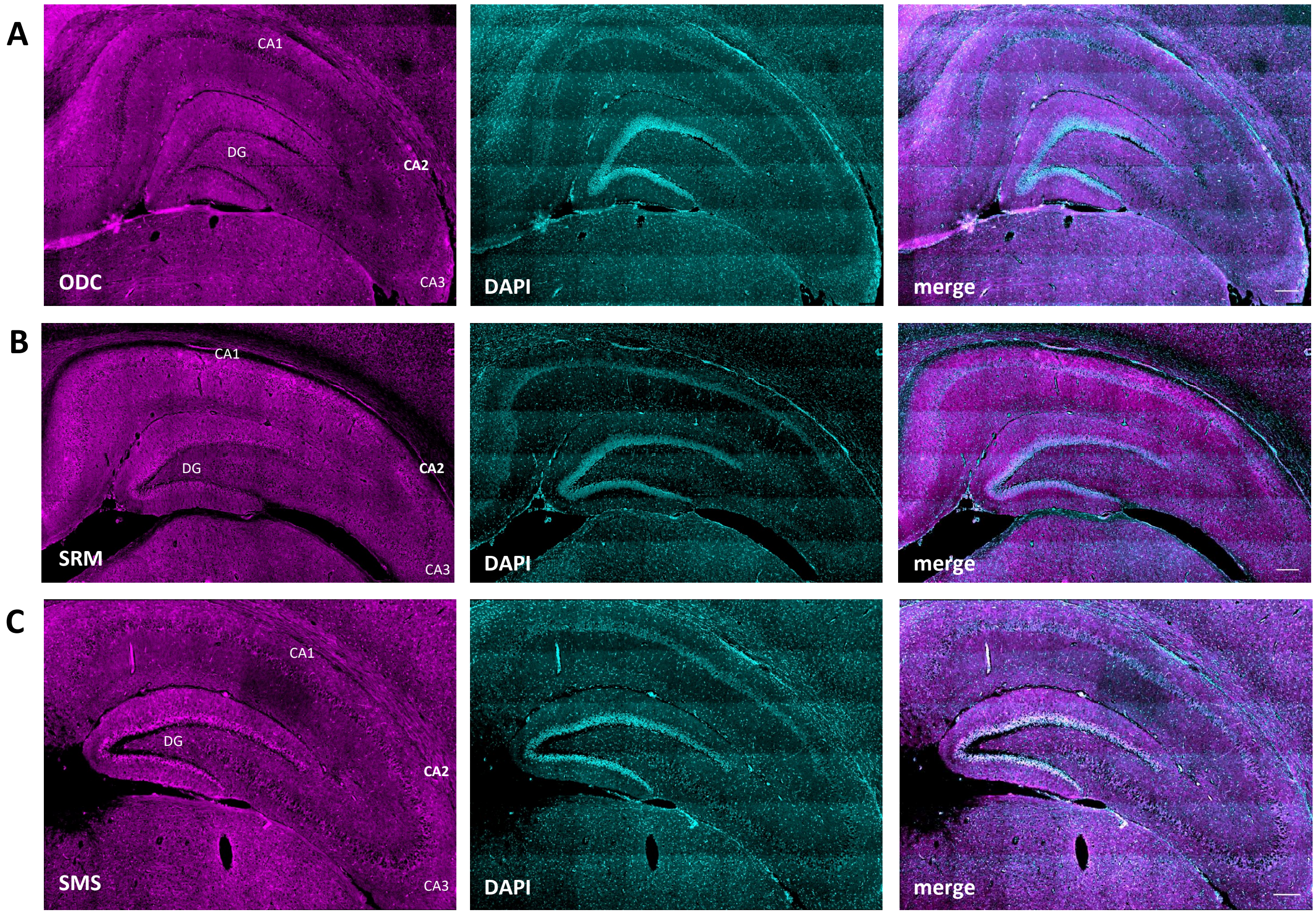


**Fig. S4. Distribution of PA synthesis-related enzymes in the rat hippocampus.** **I**mmunofluorescence staining of rat hippocampal sections showing expression of **(A)** ODC, **(B)** SRM, and **(C)** SMS (magenta). Sections were counterstained with DAPI (cyan). Panels show overview images of the entire hippocampus in separate fluorescence channels and as merged images. Representative images are shown. Scale bars: 200 µm. Abbreviations: ODC – ornithine decarboxylase; SMS – spermine synthase; SRM – spermidine synthase.

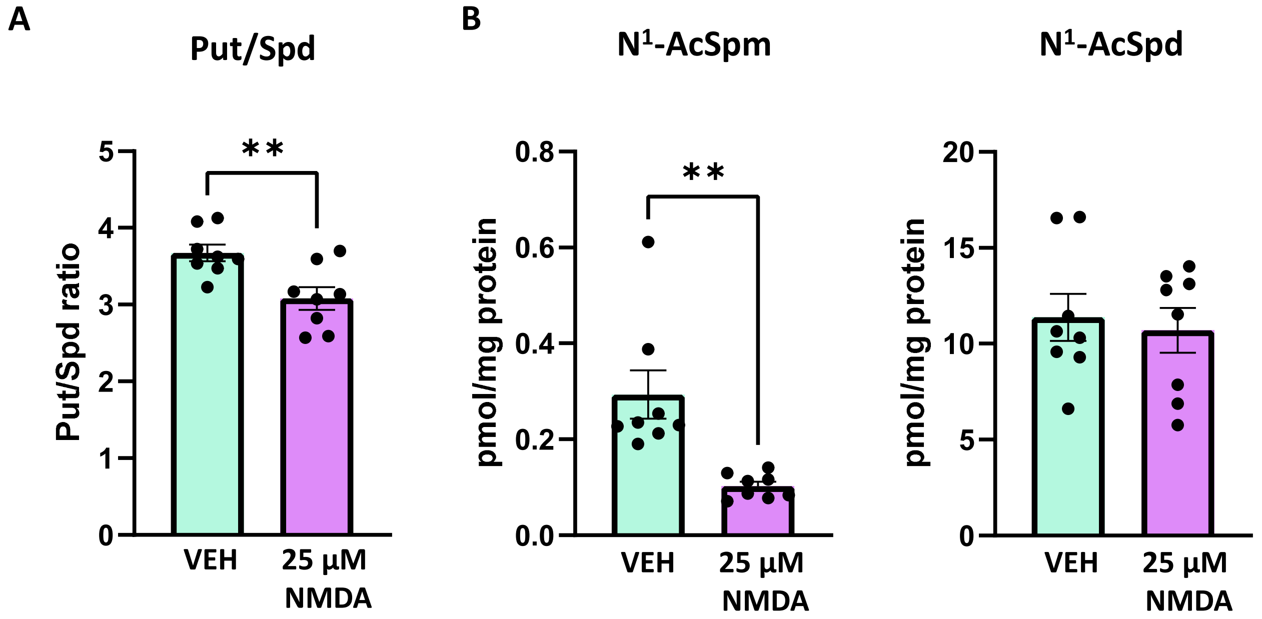


**Fig. S5. Effect of NMDA treatment on levels of Put/Spd ratio and levels of PA degradation products in the OHCs. (A)** Measurements of Put/Spd ratio in the VEH- and 25 µM NMDA-exposed OHCs. A statistically significant decrease in Put/Spd ratio was observed in NMDA-exposed OHCs compared to VEH-exposed OHCs (3.081 ± 0.15 vs. 3.673 ± 0.103, respectively; $p = 0.0058$; n = 8 biological replicates (individual cultures)). **(B)** Measurements of PA degradation products in the VEH- and 25 µM NMDA-exposed OHCs. A statistically significant decrease in N1-AcSpm levels was observed in NMDA-exposed OHCs compared to VEH-exposed OHCs (0.1025 ± 0.0090 pmol/mg protein vs. 0.2933 ± 0.0494 pmol/mg protein, respectively; $p = 0.0022$; n = 8 biological replicates (individual cultures)). No statistically significant differences in N1-AcSpd levels were observed between VEH-exposed OHCs compared to NMDA-exposed OHCs (11.38 ± 1.25 pmol/mg protein vs. 10.69 ± 1.16 pmol/mg protein, respectively; $p = 0.6949$; n = 8 biological replicates (individual cultures)). Data are presented as mean ± SEM. **$p < 0.01$, unpaired t-test. Abbreviations: $N^1$-AcSpd – $N^1$-acetyl-spermidine; $N^1$-AcSpm – $N^1$-acetyl-spermine; NMDA – n-methyl-d-aspartate; OHCs – organotypic hippocampal cultures; PA – polyamines; Put – putrescine; Spd – spermidine; VEH – vehicle.

#### Table S1. Results of Tukey's multiple comparisons test for the DFMO experiment in OHCs.

| Tukey's multiple comparisons test | Mean diff. | 95.00% CI of diff. | Summary | Adjusted P Value |
|---|---|---|---|---|
| 25µM NMDA:CA1 vs. 25µM NMDA:CA2 | 21.55 | 13,33 to 29,76 | *** | 0.0001 |
| 25µM NMDA:CA1 vs. 25µM NMDA + 0.2mM DFMO:CA1 | -2.843 | -11,06 to 5,372 | ns | 0.7956 |
| 25µM NMDA:CA1 vs. 25µM NMDA + 0.2mM DFMO:CA2 | 12.97 | 4,753 to 21,18 | ** | 0.0038 |
| 25µM NMDA:CA1 vs. 25µM NMDA + 1mM DFMO:CA1 | -3.806 | -12,02 to 4,409 | ns | 0.57 |
| 25µM NMDA:CA1 vs. 25µM NMDA + 1mM DFMO:CA2 | 12.77 | 4,559 to 20,99 | ** | 0.0041 |
| 25µM NMDA:CA2 vs. 25µM NMDA + 0.2mM DFMO:CA1 | -24.39 | -32,61 to -16,18 | **** | <0,0001 |
| 25µM NMDA:CA2 vs. 25µM NMDA + 0.2mM DFMO:CA2 | -8.581 | -16,80 to -0,3661 | * | 0.0403 |
| 25µM NMDA:CA2 vs. 25µM NMDA + 1mM DFMO:CA1 | -25.35 | -33,57 to -17,14 | **** | <0,0001 |
| 25µM NMDA:CA2 vs. 25µM NMDA + 1mM DFMO:CA2 | -8.775 | -16,99 to -0,5601 | * | 0.036 |
| 25µM NMDA + 0.2mM DFMO:CA1 vs. 25µM NMDA + 0.2mM DFMO:CA2 | 15.81 | 7,596 to 24,03 | ** | 0.001 |
| 25µM NMDA + 0.2mM DFMO:CA1 vs. 25µM NMDA + 1mM DFMO:CA1 | -0.9629 | -9,178 to 7,252 | ns | 0.9974 |
| 25µM NMDA + 0.2mM DFMO:CA1 vs. 25µM NMDA + 1mM DFMO:CA2 | 15.62 | 7,402 to 23,83 | ** | 0.0011 |
| 25µM NMDA + 0.2mM DFMO:CA2 vs. 25µM NMDA + 1mM DFMO:CA1 | -16.77 | -24,99 to -8,559 | *** | 0.0007 |
| 25µM NMDA + 0.2mM DFMO:CA2 vs. 25µM NMDA + 1mM DFMO:CA2 | -0.194 | -8,409 to 8,021 | ns | >0,9999 |
| 25µM NMDA + 1mM DFMO:CA1 vs. 25µM NMDA + 1mM DFMO:CA2 | 16.58 | 8,365 to 24,79 | *** | 0.0007 |

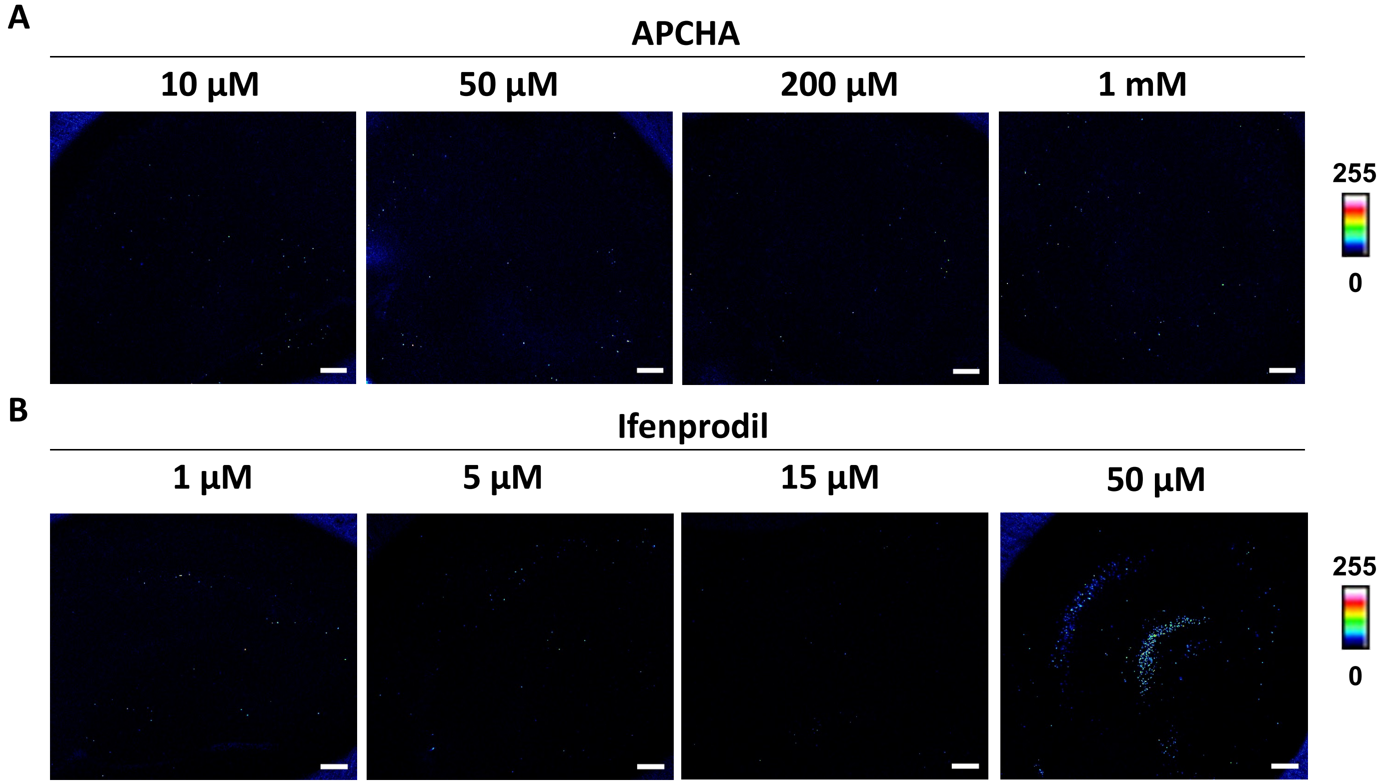


**Fig. S6. Assessment of OHC viability following exposure to different concentrations of APCHA and ifenprodil.** Representative pseudocolor fluorescence images of PI-labelled OHCs treated with increasing concentrations of **(A)** APCHA (10 µM, 50 µM, 200 µM, 1 mM) and **(B)** ifenprodil (1 µM, 5 µM, 15 µM, 50 µM). Intensity is pseudocolour-coded from 0 (low) to 255 (high). Scale bars: 200 µm. Abbreviations: APCHA – $N^1$-cyclohexylpropane-1,3-diamine; OHCs – organotypic hippocampal cultures; PI – propidium iodide.

### Table S2. Results of Tukey's multiple comparisons test for the APCHA experiment in OHCs.

| Tukey's multiple comparisons test | Mean diff. | 95.00% CI of diff. | Summary | Adjusted P Value |
|---|---|---|---|---|
| 25 µM NMDA:CA1 vs. 25 µM NMDA:CA2 | 15.48 | 7,600 to 23,35 | **** | <0,0001 |
| 25 µM NMDA:CA1 vs. 1mM APCHA + 25µM NMDA:CA1 | 5.541 | -2,336 to 13,42 | ns | 0.3292 |
| 25 µM NMDA:CA1 vs. 1mM APCHA + 25µM NMDA:CA2 | 28.61 | 20,73 to 36,48 | **** | <0,0001 |
| 25 µM NMDA:CA1 vs. 100µM SPM:CA1 | 1.179 | -6,698 to 9,055 | ns | >0,9999 |
| 25 µM NMDA:CA1 vs. 100µM SPM:CA2 | 17.67 | 9,790 to 25,54 | **** | <0,0001 |
| 25 µM NMDA:CA1 vs. 100µM SPM + 25µM NMDA:CA1 | -0.06467 | -7,941 to 7,812 | ns | >0,9999 |
| 25 µM NMDA:CA1 vs. 100µM SPM + 25µM NMDA:CA2 | 14.03 | 6,158 to 21,91 | *** | 0.0001 |
| 25 µM NMDA:CA1 vs. 1mM APCHA + 100µM SPM + 25µM NMDA:CA1 | -2.007 | -9,884 to 5,869 | ns | 0.9946 |
| 25 µM NMDA:CA1 vs. 1mM APCHA + 100µM SPM + 25µM NMDA:CA2 | 7.669 | -0,2076 to 15,55 | ns | 0.0603 |
| 25 µM NMDA:CA2 vs. 1mM APCHA + 25µM NMDA:CA1 | -9.936 | -17,81 to -2,060 | ** | 0.007 |
| 25 µM NMDA:CA2 vs. 1mM APCHA + 25µM NMDA:CA2 | 13.13 | 5,255 to 21,01 | *** | 0.0003 |
| 25 µM NMDA:CA2 vs. 100µM SPM:CA1 | -14.3 | -22,17 to -6,422 | *** | 0.0001 |
| 25 µM NMDA:CA2 vs. 100µM SPM:CA2 | 2.189 | -5,687 to 10,07 | ns | 0.99 |
| 25 µM NMDA:CA2 vs. 100µM SPM + 25µM NMDA:CA1 | -15.54 | -23,42 to -7,665 | **** | <0,0001 |
| 25 µM NMDA:CA2 vs. 100µM SPM + 25µM NMDA:CA2 | -1.443 | -9,319 to 6,434 | ns | 0.9996 |
| 25 µM NMDA:CA2 vs. 1mM APCHA + 100µM SPM + 25µM NMDA:CA1 | -17.48 | -25,36 to -9,608 | **** | <0,0001 |
| 25 µM NMDA:CA2 vs. 1mM APCHA + 100µM SPM + 25µM NMDA:CA2 | -7.808 | -15,68 to 0,06847 | ns | 0.0532 |
| 1mM APCHA + 25µM NMDA:CA1 vs. 1mM APCHA + 25µM NMDA:CA2 | 23.07 | 15,19 to 30,94 | **** | <0,0001 |
| 1mM APCHA + 25µM NMDA:CA1 vs. 100µM SPM:CA1 | -4.362 | -12,24 to 3,514 | ns | 0.6322 |
| 1mM APCHA + 25µM NMDA:CA1 vs. 100µM SPM:CA2 | 12.13 | 4,249 to 20,00 | *** | 0.0008 |
| 1mM APCHA + 25µM NMDA:CA1 vs. 100µM SPM + 25µM NMDA:CA1 | -5.605 | -13,48 to 2,271 | ns | 0.3153 |
| 1mM APCHA + 25µM NMDA:CA1 vs. 100µM SPM + 25µM NMDA:CA2 | 8.494 | 0,6171 to 16,37 | * | 0.0282 |
| 1mM APCHA + 25µM NMDA:CA1 vs. 1mM APCHA + 100µM SPM + 25µM NMDA:CA1 | -7.548 | -15,42 to 0,3283 | ns | 0.0672 |
| 1mM APCHA + 25µM NMDA:CA1 vs. 1mM APCHA + 100µM SPM + 25µM NMDA:CA2 | 2.128 | -5,748 to 10,00 | ns | 0.9918 |
| 1mM APCHA + 25µM NMDA:CA2 vs. 100µM SPM:CA1 | -27.43 | -35,31 to -19,55 | **** | <0,0001 |
| 1mM APCHA + 25µM NMDA:CA2 vs. 100µM SPM:CA2 | -10.94 | -18,82 to -3,066 | ** | 0.0026 |
| 1mM APCHA + 25µM NMDA:CA2 vs. 100µM SPM + 25µM NMDA:CA1 | -28.67 | -36,55 to -20,80 | **** | <0,0001 |
| 1mM APCHA + 25µM NMDA:CA2 vs. 100µM SPM + 25µM NMDA:CA2 | -14.57 | -22,45 to -6,698 | **** | <0,0001 |
| 1mM APCHA + 25µM NMDA:CA2 vs. 1mM APCHA + 100µM SPM + 25µM NMDA:CA1 | -30.62 | -38,49 to -22,74 | **** | <0,0001 |
| 1mM APCHA + 25µM NMDA:CA2 vs. 1mM APCHA + 100µM SPM + 25µM NMDA:CA2 | -20.94 | -28,82 to -13,06 | **** | <0,0001 |
| 100µM SPM:CA1 vs. 100µM SPM:CA2 | 16.49 | 8,611 to 24,36 | **** | <0,0001 |
| 100µM SPM:CA1 vs. 100µM SPM + 25µM NMDA:CA1 | -1.243 | -9,120 to 6,633 | ns | 0.9999 |
| 100µM SPM:CA1 vs. 100µM SPM + 25µM NMDA:CA2 | 12.86 | 4,979 to 20,73 | *** | 0.0004 |
| 100µM SPM:CA1 vs. 1mM APCHA + 100µM SPM + 25µM NMDA:CA1 | -3.186 | -11,06 to 4,690 | ns | 0.9025 |
| 100µM SPM:CA1 vs. 1mM APCHA + 100µM SPM + 25µM NMDA:CA2 | 6.49 | -1,386 to 14,37 | ns | 0.1644 |
| 100µM SPM:CA2 vs. 100µM SPM + 25µM NMDA:CA1 | -17.73 | -25,61 to -9,854 | **** | <0,0001 |
| 100µM SPM:CA2 vs. 100µM SPM + 25µM NMDA:CA2 | -3.632 | -11,51 to 4,244 | ns | 0.8176 |
| 100µM SPM:CA2 vs. 1mM APCHA + 100µM SPM + 25µM NMDA:CA1 | -19.67 | -27,55 to -11,80 | **** | <0,0001 |
| 100µM SPM:CA2 vs.1mM APCHA + 100µM SPM + 25µM NMDA:CA2 | -9.997 | -17,87 to -2,121 | ** | 0.0066 |
| 100µM SPM + 25µM NMDA:CA1 vs. 100µM SPM + 25µM NMDA:CA2 | 14.1 | 6,222 to 21,98 | *** | 0.0001 |
| 100µM SPM + 25µM NMDA:CA1 vs. 1mM APCHA + 100µM SPM + 25µM NMDA:CA1 | -1.943 | -9,819 to 5,934 | ns | 0.9957 |
| 100µM SPM + 25µM NMDA:CA1 vs. 1mM APCHA + 100µM SPM + 25µM NMDA:CA2 | 7.734 | -0,1429 to 15,61 | ns | 0.0569 |
| 100µM SPM + 25µM NMDA:CA2 vs. 1mM APCHA + 100µM SPM + 25µM NMDA:CA1 | -16.04 | -23,92 to -8,165 | **** | <0,0001 |
| 100µM SPM + 25µM NMDA:CA2 vs. 1mM APCHA + 100µM SPM + 25µM NMDA:CA2 | -6.365 | -14,24 to 1,511 | ns | 0.1813 |
| 1mM APCHA + 100µM SPM + 25µM NMDA:CA1 vs.1mM APCHA + 100µM SPM + 25µM NMDA:CA2 | 9.676 | 1,800 to 17,55 | ** | 0.009 |

**Table S3. Results of Tukey's multiple comparisons test for the ifenprodil experiment in OHCs.**

| Tukey's multiple comparisons test | Mean diff. | 95.00% CI of diff. | Summary | Adjusted P Value |
|---|---|---|---|---|
| 25 µM NMDA:CA1 vs. 25 µM NMDA:CA2 | 21.77 | 15.34 to 28.20 | **** | <0.0001 |
| 25 µM NMDA:CA1 vs. 15µM Ifenprodil + 25µM NMDA:CA1 | 11.05 | 4.623 to 17.48 | **** | <0.0001 |
| 25 µM NMDA:CA1 vs. 15µM Ifenprodil + 25µM NMDA:CA2 | 32.2 | 25.77 to 38.63 | **** | <0.0001 |
| 25 µM NMDA:CA1 vs. 100µM SPM:CA1 | 4.483 | -1.946 to 10.91 | ns | 0.4088 |
| 25 µM NMDA:CA1 vs. 100µM SPM:CA2 | 20.35 | 13.92 to 26.78 | **** | <0.0001 |
| 25 µM NMDA:CA1 vs. 100µM SPM + 25µM NMDA:CA1 | -4.604 | -11.03 to 1.825 | ns | 0.3701 |
| 25 µM NMDA:CA1 vs. 100µM SPM + 25µM NMDA:CA2 | 17.57 | 11.14 to 24.00 | **** | <0.0001 |
| 25 µM NMDA:CA1 vs. 15µM Ifenprodil + 100µM SPM:CA1 | 1.869 | -4.560 to 8.297 | ns | 0.9962 |
| 25 µM NMDA:CA1 vs. 15µM Ifenprodil + 100µM SPM:CA2 | 17.16 | 10.73 to 23.59 | **** | <0.0001 |
| 25 µM NMDA:CA1 vs. 15µM Ifenprodil + 100µM SPM + 25µM NMDA:CA1 | 0.7689 | -5.660 to 7.198 | ns | >0.9999 |
| 25 µM NMDA:CA1 vs. 15µM Ifenprodil + 100µM SPM + 25µM NMDA:CA2 | 15.93 | 9.501 to 22.36 | **** | <0.0001 |
| 25 µM NMDA:CA2 vs. 15µM Ifenprodil + 25µM NMDA:CA1 | -10.72 | -17.15 to -4.293 | **** | <0.0001 |
| 25 µM NMDA:CA2 vs. 15µM Ifenprodil + 25µM NMDA:CA2 | 10.43 | 3.996 to 16.85 | *** | 0.0001 |
| 25 µM NMDA:CA2 vs. 100µM SPM:CA1 | -17.29 | -23.72 to -10.86 | **** | <0.0001 |
| 25 µM NMDA:CA2 vs. 100µM SPM:CA2 | -1.424 | -7.853 to 5.005 | ns | 0.9997 |
| 25 µM NMDA:CA2 vs. 100µM SPM + 25µM NMDA:CA1 | -26.38 | -32.81 to -19.95 | **** | <0.0001 |
| 25 µM NMDA:CA2 vs. 100µM SPM + 25µM NMDA:CA2 | -4.207 | -10.64 to 2.222 | ns | 0.5026 |
| 25 µM NMDA:CA2 vs. 15µM Ifenprodil + 100µM SPM:CA1 | -19.9 | -26.33 to -13.48 | **** | <0.0001 |
| 25 µM NMDA:CA2 vs. 15µM Ifenprodil + 100µM SPM:CA2 | -4.617 | -11.05 to 1.812 | ns | 0.3662 |
| 25 µM NMDA:CA2 vs. 15µM Ifenprodil + 100µM SPM + 25µM NMDA:CA1 | -21 | -27.43 to -14.58 | **** | <0.0001 |
| 25 µM NMDA:CA2 vs. 15µM Ifenprodil + 100µM SPM + 25µM NMDA:CA2 | -5.843 | -12.27 to 0.5854 | ns | 0.1038 |
| 15µM Ifenprodil + 25µM NMDA:CA1 vs. 15µM Ifenprodil + 25µM NMDA:CA2 | 21.15 | 14.72 to 27.58 | **** | <0.0001 |
| 15µM Ifenprodil + 25µM NMDA:CA1 vs. 100µM SPM:CA1 | -6.569 | -13.00 to -0.1407 | * | 0.0415 |
| 15µM Ifenprodil + 25µM NMDA:CA1 vs. 100µM SPM:CA2 | 9.297 | 2.868 to 15.73 | *** | 0.0007 |
| 15µM Ifenprodil + 25µM NMDA:CA1 vs. 100µM SPM + 25µM NMDA:CA1 | -15.66 | -22.08 to -9.227 | **** | <0.0001 |
| 15µM Ifenprodil + 25µM NMDA:CA1 vs. 100µM SPM + 25µM NMDA:CA2 | 6.515 | 0.08585 to 12.94 | * | 0.0447 |
| 15µM Ifenprodil + 25µM NMDA:CA1 vs. 15µM Ifenprodil + 100µM SPM:CA1 | -9.183 | -15.61 to -2.755 | *** | 0.0009 |
| 15µM Ifenprodil + 25µM NMDA:CA1 vs. 15µM Ifenprodil + 100µM SPM:CA2 | 6.105 | -0.3240 to 12.53 | ns | 0.0755 |
| 15µM Ifenprodil + 25µM NMDA:CA1 vs. 15µM Ifenprodil + 100µM SPM + 25µM NMDA:CA1 | -10.28 | -16.71 to -3.854 | *** | 0.0001 |
| 15µM Ifenprodil + 25µM NMDA:CA1 vs. 15µM Ifenprodil + 100µM SPM + 25µM NMDA:CA2 | 4.878 | -1.551 to 11.31 | ns | 0.2903 |
| 15µM Ifenprodil + 25µM NMDA:CA2 vs. 100µM SPM:CA1 | -27.72 | -34.14 to -21.29 | **** | <0.0001 |
| 15µM Ifenprodil + 25µM NMDA:CA2 vs. 100µM SPM:CA2 | -11.85 | -18.28 to -5.420 | **** | <0.0001 |
| 15µM Ifenprodil + 25µM NMDA:CA2 vs. 100µM SPM + 25µM NMDA:CA1 | -36.8 | -43.23 to -30.37 | **** | <0.0001 |
| 15µM Ifenprodil + 25µM NMDA:CA2 vs. 100µM SPM + 25µM NMDA:CA2 | -14.63 | -21.06 to -8.203 | **** | <0.0001 |
| 15µM Ifenprodil + 25µM NMDA:CA2 vs. 15µM Ifenprodil + 100µM SPM:CA1 | -30.33 | -36.76 to -23.90 | **** | <0.0001 |
| 15µM Ifenprodil + 25µM NMDA:CA2 vs. 15µM Ifenprodil + 100µM SPM:CA2 | -15.04 | -21.47 to -8.613 | **** | <0.0001 |
| 15µM Ifenprodil + 25µM NMDA:CA2 vs. 15µM Ifenprodil + 100µM SPM + 25µM NMDA:CA1 | -31.43 | -37.86 to -25.00 | **** | <0.0001 |
| 15µM Ifenprodil + 25µM NMDA:CA2 vs. 15µM Ifenprodil + 100µM SPM + 25µM NMDA:CA2 | -16.27 | -22.70 to -9.840 | **** | <0.0001 |
| 100µM SPM:CA1 vs. 100µM SPM:CA2 | 15.87 | 9.438 to 22.30 | **** | <0.0001 |
| 100µM SPM:CA1 vs. 100µM SPM + 25µM NMDA:CA1 | -9.087 | -15.52 to -2.658 | ** | 0.001 |
| 100µM SPM:CA1 vs. 100µM SPM + 25µM NMDA:CA2 | 13.08 | 6.655 to 19.51 | **** | <0.0001 |
| 100µM SPM:CA1 vs. 15µM Ifenprodil + 100µM SPM:CA1 | -2.614 | -9.043 to 3.815 | ns | 0.951 |
| 100µM SPM:CA1 vs. 15µM Ifenprodil + 100µM SPM:CA2 | 12.67 | 6.246 to 19.10 | **** | <0.0001 |
| 100µM SPM:CA1 vs. 15µM Ifenprodil + 100µM SPM + 25µM NMDA:CA1 | -3.714 | -10.14 to 2.715 | ns | 0.6777 |
| 100µM SPM:CA1 vs. 15µM Ifenprodil + 100µM SPM + 25µM NMDA:CA2 | 11.45 | 5.019 to 17.88 | **** | <0.0001 |
| 100µM SPM:CA2 vs. 100µM SPM + 25µM NMDA:CA1 | -24.95 | -31.38 to -18.52 | **** | <0.0001 |
| 100µM SPM:CA2 vs. 100µM SPM + 25µM NMDA:CA2 | -2.783 | -9.211 to 3.646 | ns | 0.9267 |
| 100µM SPM:CA2 vs. 15µM Ifenprodil + 100µM SPM:CA1 | -18.48 | -24.91 to -12.05 | **** | <0.0001 |
| 100µM SPM:CA2 vs. 15µM Ifenprodil + 100µM SPM:CA2 | -3.192 | -9.621 to 3.236 | ns | 0.8392 |
| 100µM SPM:CA2 vs. 15µM Ifenprodil + 100µM SPM + 25µM NMDA:CA1 | -19.58 | -26.01 to -13.15 | **** | <0.0001 |
| 100µM SPM:CA2 vs. 15µM Ifenprodil + 100µM SPM + 25µM NMDA:CA2 | -4.419 | -10.85 to 2.010 | ns | 0.4297 |
| 100µM SPM + 25µM NMDA:CA1 vs. 100µM SPM + 25µM NMDA:CA2 | 22.17 | 15.74 to 28.60 | **** | <0.0001 |
| 100µM SPM + 25µM NMDA:CA1 vs. 15µM Ifenprodil + 100µM SPM:CA1 | 6.473 | 0.04392 to 12.90 | * | 0.0472 |
| 100µM SPM + 25µM NMDA:CA1 vs. 15µM Ifenprodil + 100µM SPM:CA2 | 21.76 | 15.33 to 28.19 | **** | <0.0001 |
| 100µM SPM + 25µM NMDA:CA1 vs. 15µM Ifenprodil + 100µM SPM + 25µM NMDA:CA1 | 5.373 | -1.056 to 11.80 | ns | 0.1768 |
| 100µM SPM + 25µM NMDA:CA1 vs. 15µM Ifenprodil + 100µM SPM + 25µM NMDA:CA2 | 20.53 | 14.11 to 26.96 | **** | <0.0001 |
| 100µM SPM + 25µM NMDA:CA2 vs. 15µM Ifenprodil + 100µM SPM:CA1 | -15.7 | -22.13 to -9.269 | **** | <0.0001 |
| 100µM SPM + 25µM NMDA:CA2 vs. 15µM Ifenprodil + 100µM SPM:CA2 | -0.4098 | -6.839 to 6.019 | ns | >0.9999 |
| 100µM SPM + 25µM NMDA:CA2 vs. 15µM Ifenprodil + 100µM SPM + 25µM NMDA:CA1 | -16.8 | -23.23 to -10.37 | **** | <0.0001 |
| 100µM SPM + 25µM NMDA:CA2 vs. 15µM Ifenprodil + 100µM SPM + 25µM NMDA:CA2 | -1.637 | -8.065 to 4.792 | ns | 0.9988 |
| 15µM Ifenprodil + 100µM SPM:CA1 vs. 15µM Ifenprodil + 100µM SPM:CA2 | 15.29 | 8.859 to 21.72 | **** | <0.0001 |
| 15µM Ifenprodil + 100µM SPM:CA1 vs. 15µM Ifenprodil + 100µM SPM + 25µM NMDA:CA1 | -1.1 | -7.529 to 5.329 | ns | >0.9999 |
| 15µM Ifenprodil + 100µM SPM:CA1 vs. 15µM Ifenprodil + 100µM SPM + 25µM NMDA:CA2 | 14.06 | 7.633 to 20.49 | **** | <0.0001 |
| 15µM Ifenprodil + 100µM SPM:CA2 vs. 15µM Ifenprodil + 100µM SPM + 25µM NMDA:CA1 | -16.39 | -22.82 to -9.959 | **** | <0.0001 |
| 15µM Ifenprodil + 100µM SPM:CA2 vs. 15µM Ifenprodil + 100µM SPM + 25µM NMDA:CA2 | -1.227 | -7.656 to 5.202 | ns | >0.9999 |
| 15µM Ifenprodil + 100µM SPM + 25µM NMDA:CA1 vs. 15µM Ifenprodil + 100µM SPM + 25µM NMDA:CA2 | 15.16 | 8.733 to 21.59 | **** | <0.0001 |
